# Aging trajectories of memory CD8^+^ T cells differ by their antigen specificity

**DOI:** 10.1101/2024.07.26.605197

**Authors:** Ines Sturmlechner, Abhinav Jain, Bin Hu, Rohit R. Jadhav, Wenqiang Cao, Hirohisa Okuyama, Lu Tian, Cornelia M. Weyand, Jörg J. Goronzy

**Affiliations:** Department of Immunology, Mayo Clinic, Rochester, MN 55905, USA; Robert and Arlene Kogod Center on Aging, Mayo Clinic, Rochester, MN 55905, USA; Department of Medicine, Division of Immunology and Rheumatology, Stanford University, Stanford, CA 94305, USA; Department of Medicine, Palo Alto Veterans Administration Healthcare System, Palo Alto, CA 94305, USA; Key Laboratory of Major Chronic Diseases of Nervous System of Liaoning Province, Health Sciences Institute of China Medical University, Shenyang, 110122, China; Department of Biomedical Data Science, Stanford University, Stanford, CA 94305, USA; Department of Medicine, Division of Rheumatology, Mayo Clinic, Rochester, MN 55905, USA

**Keywords:** Immune aging, memory T cells, EBV, T cell dysfunction, senescence, exhaustion, immune memory durability

## Abstract

Memory T cells are a highly dynamic and heterogeneous population that is maintained by cytokine-driven homeostatic proliferation interspersed with episodes of antigen-mediated expansion and contraction which affect their functional state and their durability. This heterogeneity complicates studies on the impact of aging on global human memory cells, specifically, it is unclear how aging drives memory T cell dysfunction. Here, we used chronic infection with Epstein-Barr virus (EBV) to assess the influence of age on memory states at the level of antigen-specific CD8^+^ T cells. We find that in young adults (<40 years), EBV-specific CD8^+^ T cells assume preferred differentiation states depending on their peptide specificity. By age >65-years, different T cell specificities had undergone largely distinct aging trajectories, which had in common a loss in adaptive and a gain in innate immunity signatures. No evidence was seen for cellular senescence or exhaustion. While naïve/stem-like EBV-specific T cells disappeared with age, T cell diversity of EBV-specific memory cells did not change or even increased. In summary, by controlling for antigen specificity we uncover age-associated shifts in gene expression and TCR diversity that have implications for optimizing vaccination strategies and adoptive T cell therapy.

## INTRODUCTION

The ability to induce immune memory is the hallmark of the adaptive immune system and the basis for one of the most successful medical interventions, the use of vaccines to induce protective immunity. While many vaccines act by generating neutralizing antibodies, mostly produced by long-lived plasma cells, a critical dimension of immune memory is the expansion and differentiation of antigen-specific memory T cells. T cell memory is best functionally defined as the ability to generate an enhanced response relative to the first encounter upon reencountering an antigen. The key desirable property of immune memory is functional durability, frequently implying a compromise between the expansion of antigen-specific memory T cells while allowing for space for T cell responses against new antigens.

One major confounding factor in memory T cell generation and homeostasis is age^1,2^. However, the immense heterogeneity of memory cells represents a challenge to define age-related mechanisms that influence the durability of functional memory T cells^3^. Cells that have encountered antigens differentiate into various memory subsets, mainly defined by their homing pattern, such as tissue-resident or recirculating central and effector memory T cells. Single cell sequencing analyses have shown that phenotypic subsetting cannot capture the full degree of transcriptional and epigenetic heterogeneity^4–6^.

A second level of complexity that complicates assessing the impact of age on memory T cell longevity comes from re-encountering antigen. Each phenotypically defined T cell subset therefore includes a composition of T cells with a range of antigen experiences, ranging from infrequent iterative boosting by reinfection or reactivation of latent viruses to continuous exposure to antigens in chronically active viral infections. Studies on memory T cell aging, even if done at the single cell level, therefore only provide estimates of averages and not age-associated trajectories, such as loss of an antigen-specific memory T cell or a change in functional states including cellular senescence or exhaustion.

Murine studies suggested that the conditions of antigen re-encounter, particularly the timing, determine the fate of peripheral antigen-specific memory T cells^7^. When antigen re-stimulation events are spaced at least 4 weeks apart, memory T cells are extraordinarily resilient to aging and persist and maintain their expansion and cytokine production potential. However, when re-stimulation intervals are short, memory T cells rapidly become dysfunctional and lose their ability to expand.

To examine the influence of age in humans on the repertoire and functional state of memory T cells in a more controlled approach, we focused on antigen-specific memory T cells against a given pathogen. We investigated the heterogeneity and aging traits of human peripheral, re-circulating memory CD8^+^ T cells specific for multiple Epstein-Barr virus (EBV) antigens. This approach allowed us to examine aging trajectories of memory T cells with differing antigen specificities that were primed simultaneously early in life.

The T cell memory response against EBV infection is a suitable model system as it leverages important characteristics of T cell heterogeneity and aging: (i) EBV infects >90% of people globally with primary EBV infections occurring predominantly before 25 years-of-age^8^. After resolution of the primary EBV infection, EBV viral DNA is retained as latent, chronic infection in memory B cells over lifetime^9^. (ii) Lymphocytes, especially CD8^+^ T cells, are vital to virus control and resolution of acute EBV infections. A subset of EBV-reactive T cells is retained as memory T cells over lifetime^10,11^. (iii) Although primed in parallel, CD8^+^ T cells specific for antigens expressed during the lytic and latent stage exhibit different trajectories in the first years after infection, both in term of frequencies and expression of CD45 isoforms^12^. (iv) Similar to other herpes viruses such as Varicella Zoster virus (VZV)^13^, EBV re-activates under immunocompromised conditions^14^ including those associated with normative aging^15^. Molecularly, EBV viral DNA is detected in the peripheral blood of adults after the age of 50 years^16,17^. While age-related reactivation remains subclinical in most older adults, EBV-related malignancies (including lymphoproliferative diseases such as diffuse large B cell lymphoma, Hodgkin lymphoma, Burkitt lymphoma and nasopharyngeal cancer) are more frequently diagnosed with advancing age^18^. (v) EBV-memory T cells are numerically largely stable into older age. Data on whether their functional traits and phenotype are sensitive to age-related dysfunction are conflicting^11,19,20^. (vi) Aging of the humoral response against EBV is less important as EBV serostatus does not change with age^8^.

By comparing EBV-specific memory CD8^+^ T cells specific for different EBV proteins, we found that they have different preferred differentiation states in young adults depending on the recognized antigen. The antigen specificity determined their fate with advancing age. While they share a trajectory of gaining innate traits at the expense of adaptive immunity signatures, the extent depended on their antigen specificity. Specifically, larger age-related changes were seen for some clones recognizing latent proteins, suggesting that these specificities account for age-related defects.

## RESULTS

### Differentiation states of EBV-specific CD8^+^ T cells in young adults

Hislop *et al*. examined the evolution of a memory response after EBV infection and concluded that the antigen-specific CD8^+^ T cell populations seen during primary infection and in the first one to ten years of asymptomatic viral persistence are different^12^. The primary response was dominated by specificities to lytic epitopes which contracted more upon resolution of the acute infection than the initially less abundant latent epitope specificities. Importantly, the nature of the antigen also determined the evolution of phenotypic changes in these populations with a greater shift to a CD45RA-positive phenotype in T cells specific to lytic antigens. To identify aging trajectories of CD8^+^ memory T cells, we decided to control for the type of recognized antigen and first perform a cross-sectional, broad phenotypic characterization of CD8^+^ T cells specific to different EBV epitopes loaded onto tetramers in a cohort of young adults (<40 years, Fig. 1A). We included 8 HLA-A*02:01^+^ individuals who had detectable EBV-specific T cells, presumably due to a clinically silent infection as they frequently occur in early childhood. We performed multiplex spectral flow cytometry with 4 EBV peptide-loaded tetramers against antigens expressed in the EBV lytic life cycle (BMLF1, BRLF1) or latent life cycle (LMP2, EBNA3C) as well as 20 cell surface markers to identify classical T cell subsets, phenotypes and cell states. We first performed a high dimensional cluster analysis of tetramer-negative, bulk CD8^+^ T cells to generate an unbiased reference map recapitulating CD8^+^ T cell subsets such as multiple clusters of naïve/stem-like cells, central memory (CM), effector memory (EM) and terminal effector memory cell with CD45RA expression (TEMRA) (Fig. 1B). To assess the phenotype and heterogeneity of EBV-specific cells (Fig. 1C), we next mapped EBV-tetramer-positive cells onto the bulk reference map and found marked differences in phenotypic states across antigen-specific T cells but also within each antigen specificity (Fig. 1D). Despite inter-individual variability, the majority of T cells against lytic antigens, BMLF1 and BRLF1, were of an end-differentiated TEMRA phenotype, while those against latent antigens, LMP2 and EBNA3C, largely lacked TEMRA cells and contained more EM cells (Fig. 1E, Supplementary Fig. 1A). Similarly, flow cytometry gating for classical surface markers signifying advanced differentiation (KLRG1^+^ CD28^−^ and CX3CR1^+^ TIGIT^+^) found a higher proportion of end-differentiated cells in EBV lytic-specific T cells than in EBV latent-specific T cells (Supplementary Fig. 1B-D). To statistically compare the distribution of antigen-specific cells in these high-dimensional clustering data, we performed linear regression on probability vectors summarizing the cluster-specific cell frequencies. We found that T cells against lytic antigens were more similar to each other, while those against latent antigens, especially LMP2, differed significantly (Fig. 1F). This phenotypic divergence between EBV lytic and latent-specific T cells was also underscoredby principal component analysis (PCA) based on T cell cluster distribution data (Fig. 1G). Since the T cell differentiation stage is tightly associated with the expression of differentiation-associated transcription factors (TFs), we perform flow cytometry assessments with EBV tetramers and intracellular antibody staining. Indeed, we found that EBV lytic-antigen specific T cells expressed higher T-BET and RUNX3 levels and contained more cytotoxic effector molecules, perforin and granzyme B (Fig. 1H, Supplementary Fig. 1E). However, other TFs that correlated with T cell differentiation, such as TOX/TOX2, did not show clear differences between lytic and latent-antigen specific T cells suggesting a more complex picture beyond divergence of EBV lytic and latent antigen-specific T cells.

**Figure 1:**
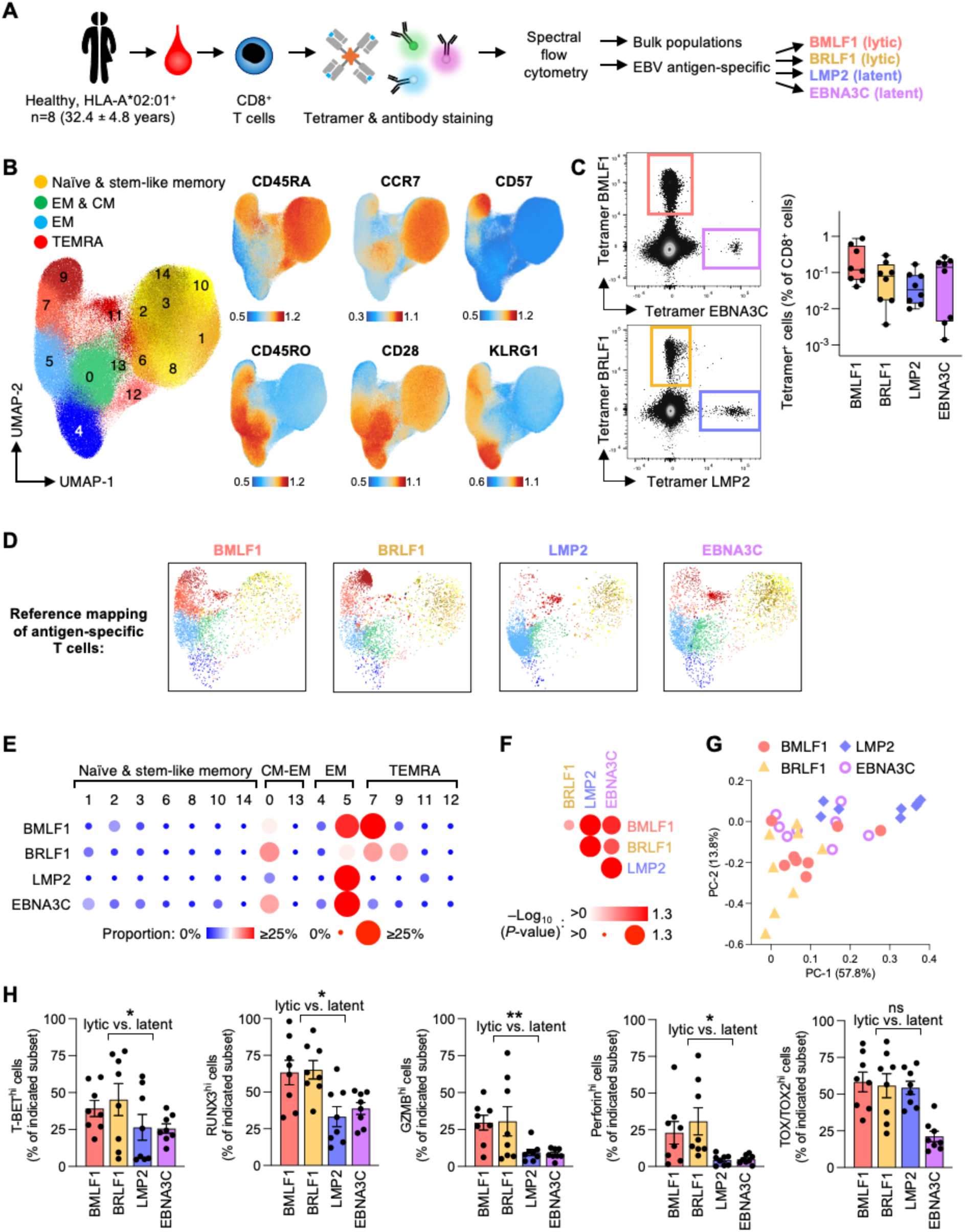
EBV-specific CD8^+^ T cells differ in their preferred, phenotypically defined differentiation states dependent on the recognized antigen. A. Schematic of experimental design: CD8^+^ T cells from peripheral blood of young adults (<40 years) were stained with the panel of antibodies shown in Supplementary Table 2 and HLA-A*02:01 tetramers loaded with indicated EBV peptides and then subjected to spectral flow cytometry. B. UMAP reference plots and feature plots of indicated cell surface marker showing bulk CD8^+^ T cell subsets. Functional subset assignment based on classical marker profiles including CD45RA/RO, CCR7, and CD28. C. Representative flow cytometry scatter plots of tetramer staining (left) and box plot of tetramer-positive cell frequencies (right). Fluorophores for tetramers were PE-Cy5 (BMLF1), BV480 (EBNA3C), APC (BRLF1), and PE (LMP2). D. Reference mapping of tetramer-positive cells projected on UMAP plots from bulk CD8^+^ T cells. E. Bubble chart of tetramer-positive cell distributions among bulk CD8^+^ T cell-defined clusters defined in Fig. 1B. F. Comparison of probability vectors reflecting the subset distributions of tetramer-positive cells. Significance levels are shown as bubble size and color. G. Principal component analysis (PCA) of cell frequency distributions. H. Protein levels of indicated transcription factors or cytotoxic molecules in tetramer-positive cells as assessed by spectral flow cytometry. Data show the median (C, E) or mean ± SEM in (H). All datapoints represent distinct biological replicates. Data were compared by two-tailed, unpaired *t*-tests (H). *P<0.05, **P<0.01. ns, not significant.

To support our targeted flow cytometry assessments, we leveraged a published single cell-sequencing dataset containing antigen-specific CD8^+^ T cells of predominantly young adults^21^. We extracted data for EBV-specific T cells against peptides from 8 different EBV proteins (lytic antigens BMLF1, BRLF1, BZLF1, and latent antigens LMP1, LMP2/2A, EBNA3A, Supplementary Fig. 2A). Cluster analysis based on targeted transcriptome and surface proteome data (Supplementary Fig. 2B-F) confirmed the preference of lytic EBV antigen-specific T cells to acquire an end-differentiated TEMRA phenotype, while latent EBV-specific T cells lacked a TEMRA contribution and contained more CM and EM cells. These differences not only group lytic versus latent EBV-specific T cells but also indicate distinctions of each antigen specificity (Supplementary Fig. 2D-E). As such, these data suggest that each antigen-specific T cell population harbors unique phenotypic and transcriptional traits despite undergoing T cell priming simultaneously early in life.

### Epigenetic profiling distinguishes signatures for each EBV-specific T cell population

ATAC-seq analysis of the open chromatin landscape is a powerful tool to profile the epigenetic state even of rare, antigen-specific T cells^22,23^. We performed ATAC-seq on fresh, FACS-purified EBV-specific CD8^+^ T cells against BMLF1, BRLF1 and LMP2 as well as sorted bulk populations of classical CM (CD45RA^−^ CCR7^+^), EM (CD45RA^−^ CCR7^+^) and TEMRA (CD45RA^−^ CD28^−^) cells (Supplementary Fig. 3A-B). We included naïve CD8^+^ T cell data from our previously published ATAC-seq dataset^24^. While classical bulk subsets demonstrated a gradual PC1 increase correalated with progressive differentiation, lytic BMLF1-specific cells showed TEMRA-like preference and latent LMP2-specific cells CM-like clustering (Fig. 2A). Although being a lytic EBV antigen, BRLF1-specific cells showed broad clustering across PC1 suggesting a more amendable epigenetic fate and a less committed cell state. K-means clustering of ChromVar scores signifying inferred TF accessibility in ATAC-seq peaks^25^ (Supplementary Fig. 3C-D) as well as K-means clustering on differential peaks (Fig. 2B) confirmed the phenotypic dominance for BMLF1 as TEMRA- and EM-like and LMP2 as CM-like. Again, BRLF1-specific cell profiles were least constrained and showed signatures of both CM-like cells as well as TEMRA- and EM-like features. TF motif enrichment analysis suggested concerted actions of key TFs to promote T cell differentiation states and features in each K-means cluster (Fig. 2B, Supplementary Fig. 4A). For each cluster, we identified genes associated with nearby promoter and intragenic peaks and determined their putative biological contributions. Cluster 1 peaks were shared between T cells specific to lytic EBV antigen BMLF1 and BRLF1 cells; functional attributes of their associated genes included cytotoxicity, cytokine signaling and TCR signaling as well as NK cell-like and innate immune features (Fig. 2C). Consistent with our flow cytometry analyses (Fig. 1), chromatin at the perforin (*PRF1*), granzyme B (*GZMB*) and *CX3CR1* loci was more open in lytic EBV antigen-specific T cells. *ZEB2*, a master TF of CD8^+^ T cell terminal differentiation as well as NK maturation^26–28^, was the gene with the most differentially open peaks in K-means cluster 1 and may orchestrate many of these traits. K-means cluster 3 was most specific to lytic BMLF1-specific cells and may reflect higher TCR signaling and cytokine production, including *IFNG* (Supplementary Fig. 4B). Cluster 4 (CM-like signature) and cluster 5 (stem-like signature) were dominant for latent LMP2-specific cells and was enriched for pathways fundamental to T cell homeostasis including *LEF1*, WNT^29^ and mTOR^30^ signaling, as well as features of T-helper differentiation (Fig. 2D, Supplementary Fig. 4C). These features suggest that LMP2-specific cells are not only less differentiated but have a higher self-renewal capacity and homeostatic fitness. K-means cluster 7 was most similar between lytic BRLF1-specific and EM/TEMRA bulk cells and emphasized enhanced cytotoxicity, cytokine production and innate immune features via a different set of genes, including the top gene HELIOS (*IKZF2*) that fine-tunes T cell chromatin remodeling, differentiation and function^31–33^ (Supplementary Fig. 4D). Collectively, these data extend our observation on phenotypic and transcriptomic heterogeneity and show that T cell populations specific for distinct EBV antigens also differ at the epigenetic level. This finding underscores the critical need to investigate memory T cell fate decisions and aging trajectories at an antigen-specific and single-cell level.

**Figure 2:**
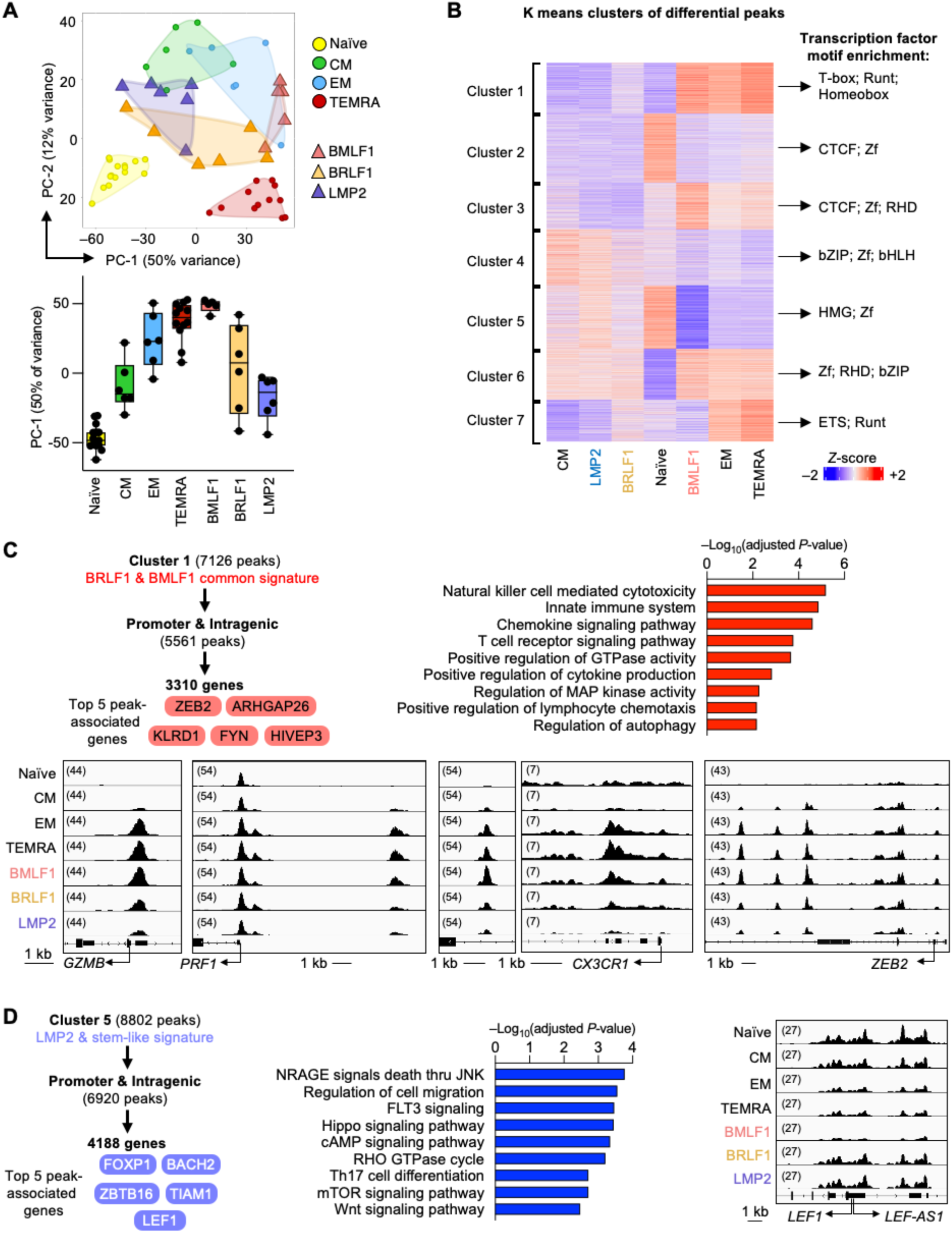
EBV-specific T cells of differing antigen specificities are distinguished in their chromatin accessibility landscapes. A. PCA of ATAC-seq data of CD8^+^ bulk subsets and tetramer-positive, EBV-specific CD8^+^ T cells. PCA is based on 5000 most variable accessible sites across all groups. PC-1 vs PC-2 scatter plot (top) and box plot of PC-1 (bottom). **B**. K-means clustering of sites differentially accessible in any binary comparison across all groups. Peaks of each cluster were analyzed for transcription factor motif enrichment via HOMER. Key transcription factor families of enriched motifs are shown at the right margin. **C-D**. Genes corresponding to peaks within each K-means cluster were identified via ChIPseeker. Only promoter and intergenic peak-associated genes were used to depict the top 5 peak-associated genes (as defined having the highest number of differential peaks in the corresponding K-means cluster) as well as for pathway and annotation enrichment. Selected ATAC-seq peak tracks are shown for K-means cluster 1 (C) and 5 (D). Data show the median (A, boxplot).

### Each antigen-specific T cell population exhibits a distinct differentiation and aging signature

To examine memory T cell aging while accounting for antigen specificity and heterogeneity, we first compared the phenotypic adaptations of EBV-specific CD8^+^ T cells in young (<40 years) and older adults (>65 years) by tetramer staining and multiplex spectral flow cytometry. As expected, naïve and stem-like cells in bulk CD8^+^ T cell population were markedly reduced with age, while end-differentiated TEMRA cells accumulated (Fig. 3A, Supplementary Fig. 5A-C). As previously described^11,34^, none of the EBV-specific T cell populations changed significantly in overall frequencies (Fig. 3B). A subset of antigen-specific T cells, especially against latent EBNA3C, changed their phenotype. Unbiased reference mapping demonstrated that EBNA3C-specific T cells gained a TEMRA population with advanced age, while the latent LMP2-specific T cells largely retained their dominant EM phenotype across age (Fig. 3C-E, Supplementary Fig. 5D). Lytic BMLF1- and BRLF1-specific cells stayed consistently more differentiated and TEMRA-like in young and older adults. Targeted flow cytometry gating on classical end-differentiation markers, TFs and effector molecules, underscored the age-related acquisition of end differentiation traits selectively in latent EBNA3C-specific cells (Fig. 3F, Supplementary Fig. 5E). Statistical analyses by comparing probablity vectors of each antigen specificity reached significance (Fig. 3E). Collectively, these data suggest that only a subset of T cells are sensitive to phenotypic adaptation with age, while other populations are capable of effectively maintaining their phenotype and avoiding age-related adaptation towards TEMRA cells.

**Figure 3:**
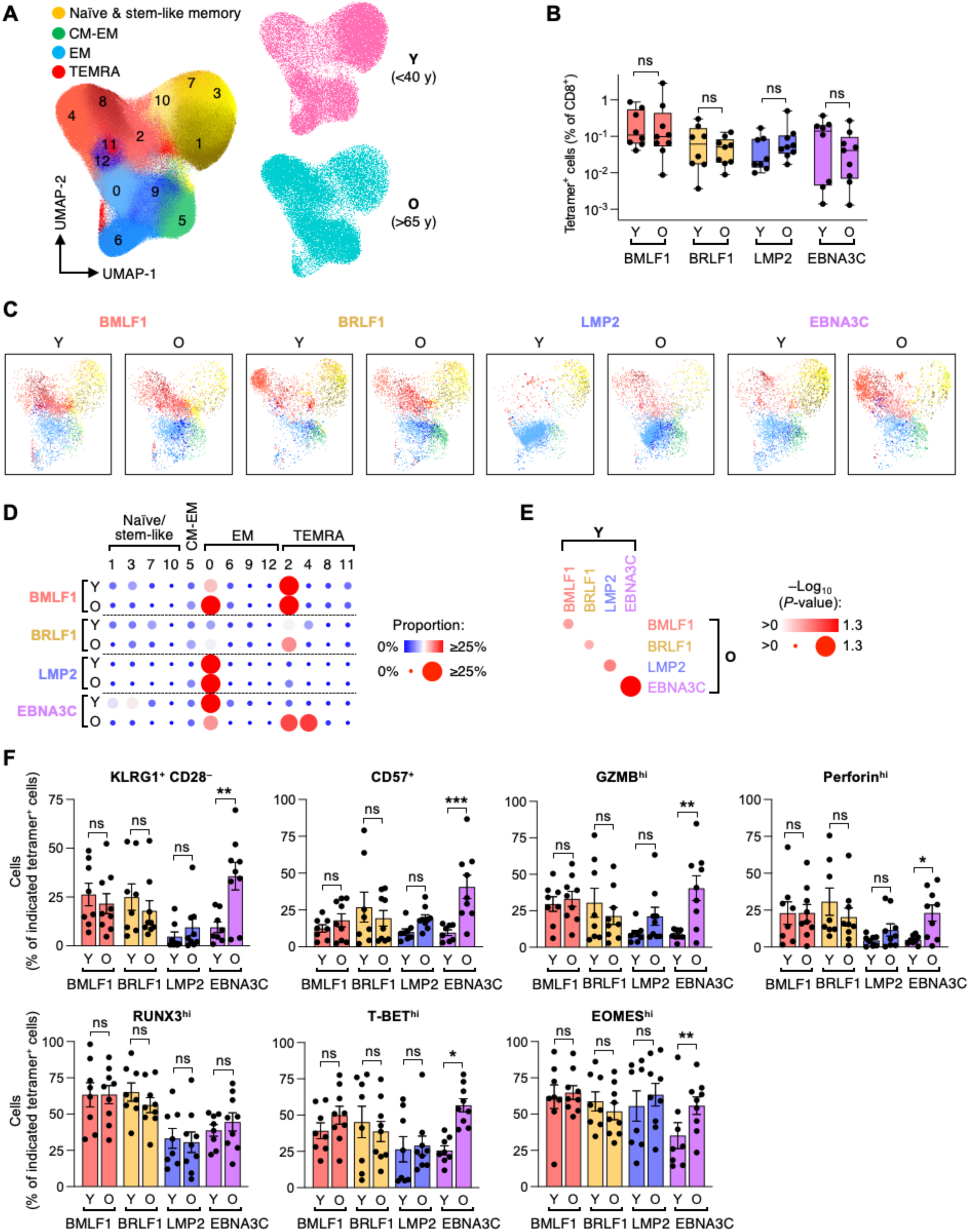
Antigen-specificity correlates with age-related phenotypic changes. CD8^+^ T cells from peripheral blood of 9 older (O) adults were analyzed by spectral flow cytometry as described in Fig. 1 and contrasted to those of 8 young (Y) adults from Fig. 1 and Supplementary Fig. 1. **A**. UMAP reference plot of bulk CD8^+^ T cell subsets based on integrated data from Y and O adults (left); distributions shown separately for Y and O (right). **B**. Frequencies of EBV tetramer-positive cells for different antigens. **C**. Reference mapping results depicting side-to-side comparison of the phenotypic distribution of tetramer-positive cells in Y and O adults. **D**. Bubble plot comparing distributions of tetramer-positive cells among CD8^+^ T cell clusters defined in (A). **E**. Probability vectors describing the subset distributions of tetramer-positive cells in Y and O adults were compared. Significance levels are shown as bubble size and color gradient. **F**. Protein levels of indicated molecules in tetramer-positive cells of Y and O adults as assessed by spectral flow cytometry. Data show the median (B, D) or mean ± SEM (F). All datapoints represent distinct biological replicates. Data were compared by two-way ANOVA with Šídák’s multiple comparisons test (B, F). Probability vectors were compared by linear regression analysis (E). *P<0.05, **P<0.01, ***P<0.001. ns, not significant.

### Single cell-sequencing with BEAM-T captures the heterogeneity of antigen-specific cells

To gain a comprehensive and unbiased snapshot of aging trajectories of antigen-specific T cells, we performed single cell-sequencing of CD8^+^ T cells from 7 young (<40 years) and 7 older adults (>65 years) with 4 modalities. We leveraged a novel method, Barcode Enabled Antigen Mapping of T cells (BEAM-T), to capture antigen specificity in addition to unbiased gene expression data, as well as surface proteome staining and TCR sequences. For each HLA-A*02:01-positive individual we extended our analysis to additional antigen-specific populations and collected BEAM-T-positive T cells specific for 4 lytic EBV antigens (BMLF1, BRLF1, BMRF1, BALF1), 4 latent EBV antigens (LMP1, LMP2, EBNA1, EBNA3C) and 2 antigens (IE62, IE63) from the related varicella zoster virus (VZV; Fig. 4A). CD8^+^ bulk T cells were collected for comparison. We obtained data on 29 304 antigen-specific T cells that passed the quality control criteria. BALF1 and IE63 yielded too few cells for downstream analyses. Antigen-specific T cells were clustered into 17 clusters representing 8 broad, classical T cell subsets (Supplementary Fig. 6A-B) and including rare T cell types with NKT-like and MAIT phenotypes (Supplementary Fig. 6B-C).

**Figure 4:**
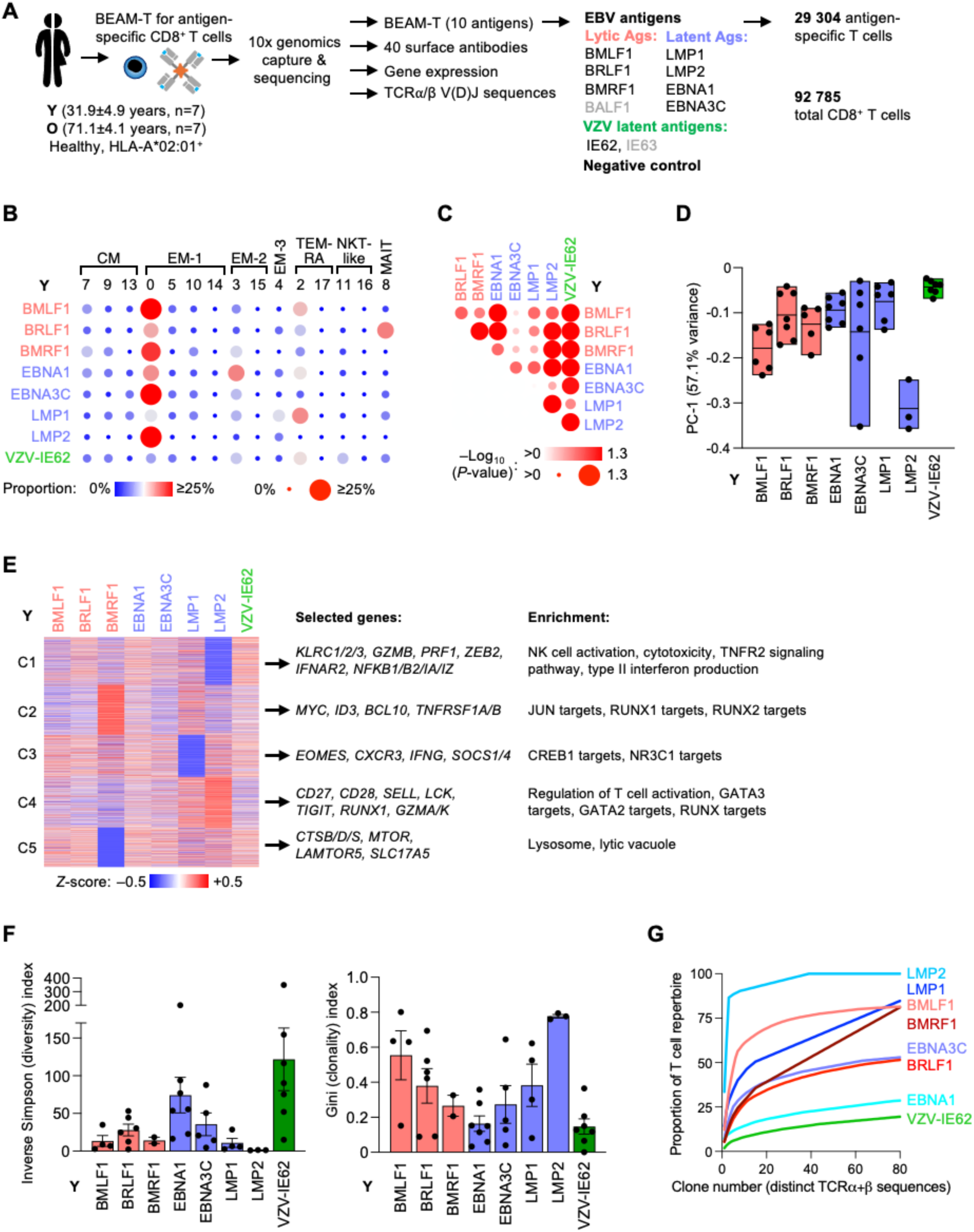
Transcriptional profiles and clonal diversity of EBV-specific T cells in young adults depend on antigen-specificity. A. Schematic of single cell RNA-seq studies of antigen-specific CD8^+^ T cells from 7 young (Y) and 7 older (O) adults. BEAM-T (Barcode Enabled Antigen Mapping of T cells) was employed. **B**. Bubble plot of antigen-specific T cell proportions among clusters defined in Supplementary Fig. 6A. **C**. Probability vectors describing the subset distribution of antigen-specific T cells were compared. Significance levels are shown as bubble size and color gradient. **D**. PCA of cell subset frequencies (see Supplementary Fig. 6F), PC-1 is shown as box plot. **E**. K-means clustering of differentially expressed genes in any binary pseudobulk comparison of antigen-specific memory T cell populations. Pathway and annotation enrichment in each K-means cluster and selected genes are shown on the right. **F**. T cell receptor (TCR) diversity and clonality are shown as Inverse Simpson (left) and Gini index (right), respectively. Only cells with TCRα and TCRβ sequences in single cell-sequencing data were used for the calculation. **G**. Cumulative TCR clone numbers depicting the numbers of distinct antigen-reactive TCR chains ordered by decreasing clonal sizes plotted against the cumulative space they occupy. Line graphs with different colors illustrate the clonal size distributions for different antigen specificities. Data show the median (B), mean (D) or mean ± SEM (F). All datapoints represent distinct biological replicates. Probability vectors were compared by linear regression analysis (C).

First, we assessed the heterogeneity of antigen-specific T cells in young adults. All populations except for latent LMP2 contained a large proportion of naïve/stem-like cells (Supplementary Fig. 6D). In line with our flow cytometry data, antigen-specific populations were heterogenous in their phenotype, but each antigen-specific T cell population exhibited their own unique subset distribution that with few exceptions was significantly different to other antigen-specific populations (Fig. 4C-D, Supplementary Fig. 6F). K-means clustering of differentially expressed transcripts followed by pathway and annotation enrichment analyses corroborated the distinct feature of each antigen-specific population (Fig. 4E). For example, unlike all other populations, latent LMP2-specific cells were devoid of NK-like and cytotoxic genes, while these cells showed the highest expression of co-activating receptors *CD27* and *CD28*.

When assessing TCRα/μ sequences in our single cell-sequencing data, we found that diversity (Inverse Simpson index or Shannon index) as well as clonality (Gini index) of antigen-specific cells varied across a broad range that did not correlate with the classification into lytic and latent antigen (Fig. 4F, Supplementary Figure 6G) and was irrespective of their dominant phenotype. This contrasts with the bulk level where the end-differentiated TEMRA phenotype is typically associated with high clonality and lower T cell diversity^35^. In fact, the lowest diversity was seen for latent LMP2 cells that were devoid of TEMRA and had predominantly acquired an EM phenotype (Fig. 4F). Conversely, cells specific for VZV-IE62, an antigen expressed in the lytic and latent VZV life cycles, had the highest TCR diversity. These data show that antigen-specificity more than differentiation state influences repertoire diversity in young individuals.

### Antigen-specific naïve and stem-like T cells are lost with aging

Next, we interrogated aging trajectories with consideration of three levels: phenotype, transcriptional profiles, and clonal diversity. The most striking age-related phenotypic change was the loss of naïve/stem-like T cells across most antigen-specific T cells (Fig. 5A), which accounted for a separation of young and older samples in PCA based on subset frequencies (Fig. 5B). A PCA focused on only the memory T cell compartment showed age-related differentiation state shifts that varied with the antigen (Fig. 5C). Specifically, latent antigen-specific T cell populations that had a low TEMRA contribution in young adults (especially EBNA1 and EBNA3C), accumulated this end-differentiated cell type in older age (Fig. 5D, Supplementary Fig. 7A) resulting in significantly different probability vectors (Fig, 5E). Lytic EBV-specific memory T cells were phenotypically more stable than those against latent antigens, with the exception of BMRF1 that frequently were MAIT-like cells in young adults that were lost with age. Latent LMP2-specific cells were an exception, they maintained their phenotypic state surprisingly well over lifetime.

**Figure 5:**
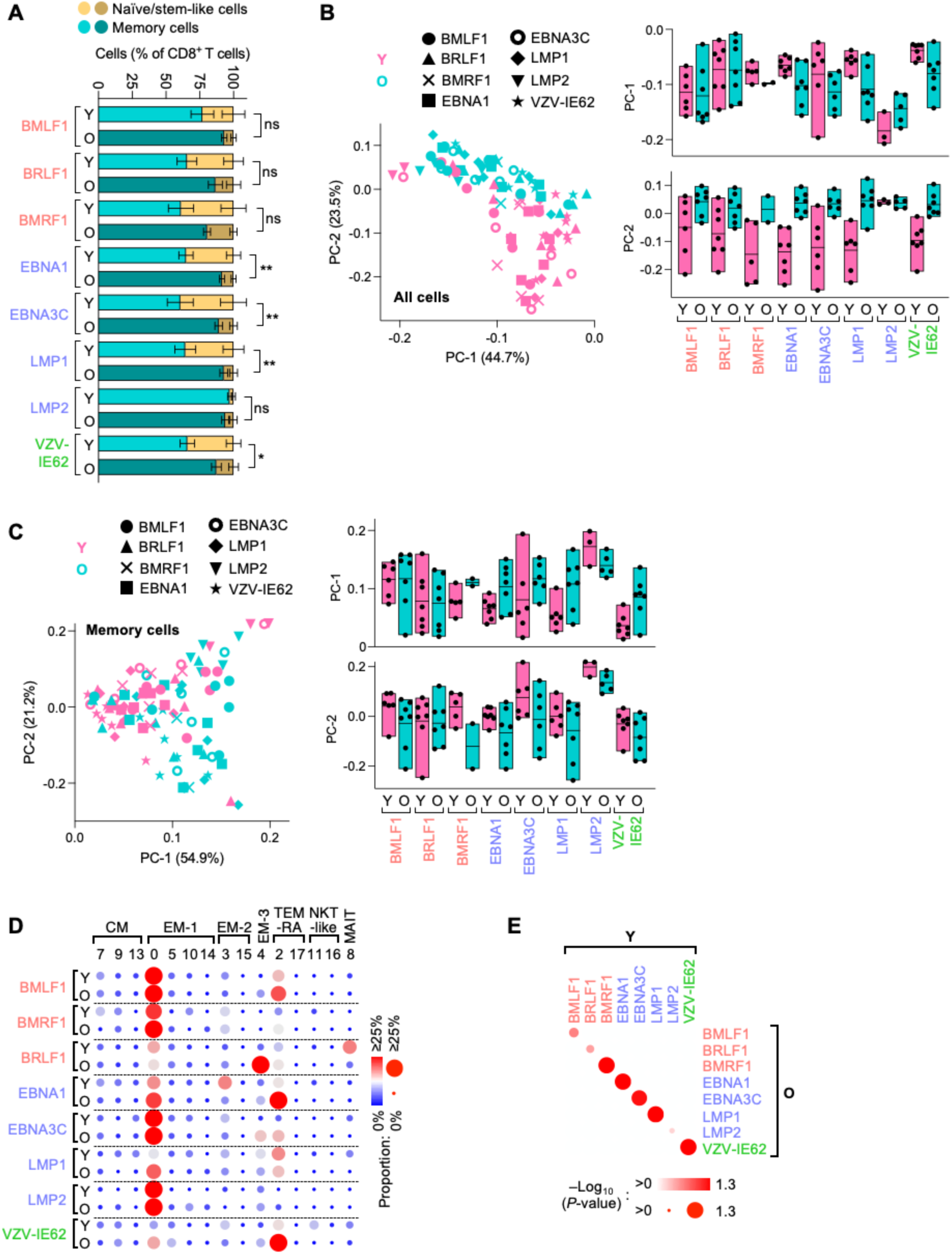
Aging results in a loss of the T cell reserve of undifferentiated antigen-specific T cells across most antigens. Single cell-sequencing of antigen-specific CD8^+^ T cells from peripheral blood of 7 young (Y) and 7 older (O) adults as described in Fig. 4 and Supplementary Fig. 6. **A**. Proportions of naïve/stem-like and memory cells, as defined in Supplementary Fig. 6A, in different antigen-specific T cell populations. **B**. PCA based on subset frequency distributions of antigen-specific cells across all clusters including naïve/stem-like cells (left). Data are shown as scatter plot (right) or box plots of PC-1 and PC-2. **C**. PCA as in (B) but based on subset frequency distributions of antigen-specific cells among memory T cell clusters after exclusion of antigen-specific cells expressing a naïve phenotype based on transcriptome or cell surface markers (Supplementary Fig. S6A-B). **D**. Bubble plot depicting distributions of antigen specific-cells among T cell subset clusters defined in Supplementary Fig. 6A. **E**. Comparison of probability vectors describing T cell subset distributions of different antigen-specific T cells. Y versus O comparisons were made only within each antigen. Significance levels are shown as bubble size and color. Data show the mean ± SEM (A) or median (B, C, D). All datapoints represent distinct biological replicates. Probability vectors were compared by linear regression analysis (E).

### Transcriptional signatures distinguish aging trajectories of memory T cells

Antigen-dependent adaptations in T cell differentiation states with age were even more apparent when we assessed transcriptional trajectories. We compared memory T cells of young and older adults for each antigen-specific population by pseudobulk differential expression analyses and found highly variable amounts of differentially expressed genes (DEGs), between 45 and 701 genes, depending on the antigen. Latent EBV-specific T cells typically exhibited higher numbers of DEGs, consistent with their higher sensitivity towards phenotypic change with age (Supplementary Fig. 7B). However, although latent LMP2-specific cells appeared stable in their cell differentiation stage, these cells had the most transcriptional changes, including many unique DEGs (Fig. 6A), suggesting qualitative changes not reflected by T cell differentiation. Consistent with the notion of population-specific aging trajectories, we find minimal overlap of DEGs across cells with different specificities (Fig. 6B). Sets of DEGs tended to be detected in cells specific to similar types of antigens, such as the nuclear antigens EBNA1 and EBNA3C, but even there they were small.

**Figure 6:**
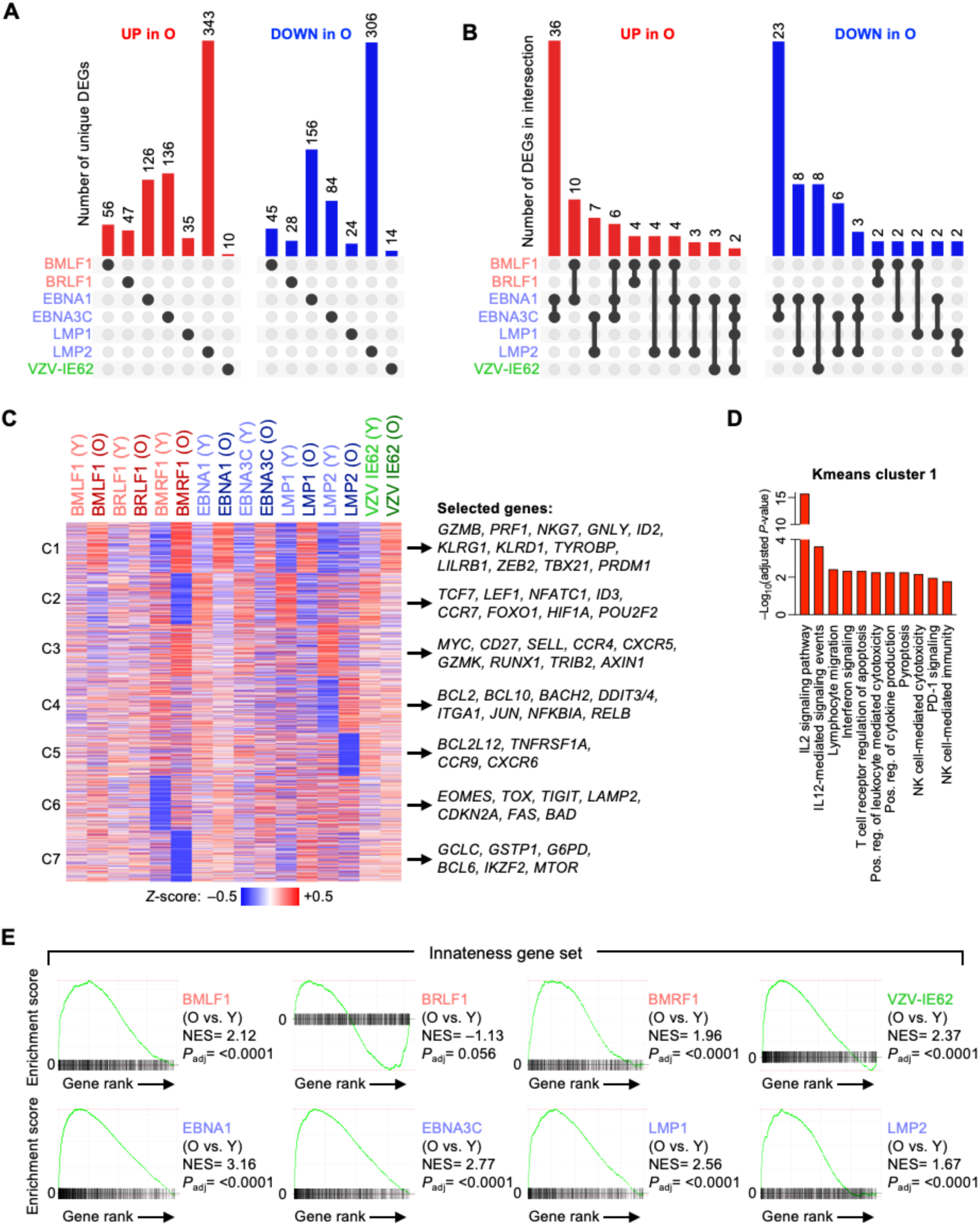
Gain of innateness in T cells is a common age-associated transcriptional signature. Pseudobulk differential expression of single cell-sequencing data from young (Y) and older (O) adults within each antigen specificity were compared and differentially expressed genes (DEGs) were identified (see also Supplementary Fig. 7B). **A-B**. Number of unique (A) and shared (B) DEGs. **C**. Heatmap and K-means clustering of DEGs identified in Supplementary Fig. 7B. Selected genes in each K-means cluster are listed on the right. **D**. Pathway and annotation enrichment analysis of DEGs in K-means cluster 1. For enrichment analyses of other K-means clusters, see Supplementary Fig. 7C. No enrichments were found for K-means clusters 5 and 6. **E**. Gene set enrichment analyses in single cell-sequencing data based on pseudobulk expression analyses of antigen-specific memory T cells comparing Y and O groups. The innateness gene set was initially collated by Gutierrez-Arcelus *et al*.^36^. NES, normalized enrichment score.

To uncover transcriptional patterns common or specific to antigen-specific T cell populations, we performed K-means clustering on all DEGs (Fig. 6C). Of the 7 clusters, clusters 1 (C1) and 2 (C2) were particularly informative as they largely represented common aging signatures. C1 genes increased with age and included cytotoxicity genes (*GZMB*, *PRF1*, *NKG7*), advanced differentiation marker (*KLRG1*, *LILRB1*/ CD85J) and genes typically expressed in NK cells (*KLRD1*, *TYROBP*). Pathway and annotation enrichment on C1 genes substantiated that a gain in cytotoxicity, NK cell-like features as well as cytokine and TCR signaling were common traits of antigen-specific memory T cell aging (Fig. 6D). C2 genes were often reduced with age and included *TCF1*, *LEF1*, *ID3*, *CCR7* that are part of essential pathways promoting T cell longevity, self-renewal capacity or lymph node homing properties, although enrichment for these pathways did not reach significance (Supplementary Fig. 7C). Antigen-specific T cells against VZV-IE62 showed similar age-related transcriptional patterns suggesting conserved features of antigen-specific memory T cell aging, at least across chronic viral infection models. Gene expression of latent EBV LMP2-specific T cells was a notable exception showing no change in C1 or C2 genes, but instead favored C3, C4 and C5 clusters. Interrogation of the respective DEGs and enriched pathways suggested an age-related increase in stress-response and survival capacity (*BCL2*, *DDIT3*, *DDIT4*) as well as pro-inflammatory NFκB signaling (*RELB*, *NFKBIA*). However, these cells also lost features of T cell longevity (*MYC*, *RUNX1*, *CD27*, *SELL*/CD62L) suggesting complex adaptations in memory T cell fitness.

Loss of essential T cell fidelity genes (C2) and gain in NK cell traits (C1) suggested a rewiring of adaptive and innate immunity programs. Specifically, cytotoxic features have been previously correlated with innateness, while *TCF7*, *MYC* and other key TFs have been correlated with T cell adaptiveness^36^. To test this hypothesis, we performed gene set enrichment analyses (GSEA) for gene modules previously defined as hallmarks for innateness or adaptiveness^36^. We found a highly significant, positive enrichment for innateness genes in 7 out of 8 antigen-specificities in older adults (Fig. 6E) indicating that a gain in innateness is a common signature of memory T cell aging. Conversely, the adaptiveness signature was lost with aging in 6 out of the 8 antigen-specific T cell populations (Supplementary Fig. 7D) indicating a transcriptional, age-sensitive switch towards innate T cell functions such as those seen in ψο T cells and NK cells. In contrast, GSEA for other, classical states of cellular dysfunction such as T cell exhaustion or cellular senescence were not significantly enriched in EBV-specific T cells of older adults (Supplementary Fig. 7E).

### Antigen-specific T cells exhibit distinct propensities towards repertoire contraction versus diversification with advancing age

Aging reduces T cell diversity of the CD8^+^ T cell compartment predominantly due to a loss of CD8^+^ T cells with a naïve/stem-like phenotype, while increasing clonality^37^. Consistent with this observation in bulk T cells, the naïve/stem-like T cell pool of antigen-specific T cells that still existed in young adults was essentially lost accounting for a contraction in diversity (Fig. 7A). In contrast, TCR diversity of antigen-specific T cells expressing memory markers was largely unchanged with advancing age for most specificities (Fig. 7B). Clonality as determined by the Gini index was increased in 2 (EBNA1, VZV-IE62) of the 6 assessed antigen-specific populations, while unchanged or even reduced with age for the other antigens (Fig. 7C). Similarly, the fraction of large T cell clones was only increased for these same 2 specificities (Fig. 7D). For a more intuitive illustration of T cell diversity, we blotted clone numbers against the cumulative space they occupy. Increased clonality with age, i.e., fewer clones occupying a given space, was seen for the responses to the latent antigens VZV-IE62, EBNA1 and EBNA3C. In contrast, a repertoire diversification was seen for the antigens BMLF1, BRLF1, and LMP1 (Fig. 7E), possibly indicating a recruitment of naïve/stem-like cells into the memory T cell compartment over adult lifetime. Taken together, aging depleted the resource of single antigen-specific naïve/stem-like cells while repertoire diversity of antigen-specific effector memory cells did not change much or even increased depending on the antigen.

**Figure 7:**
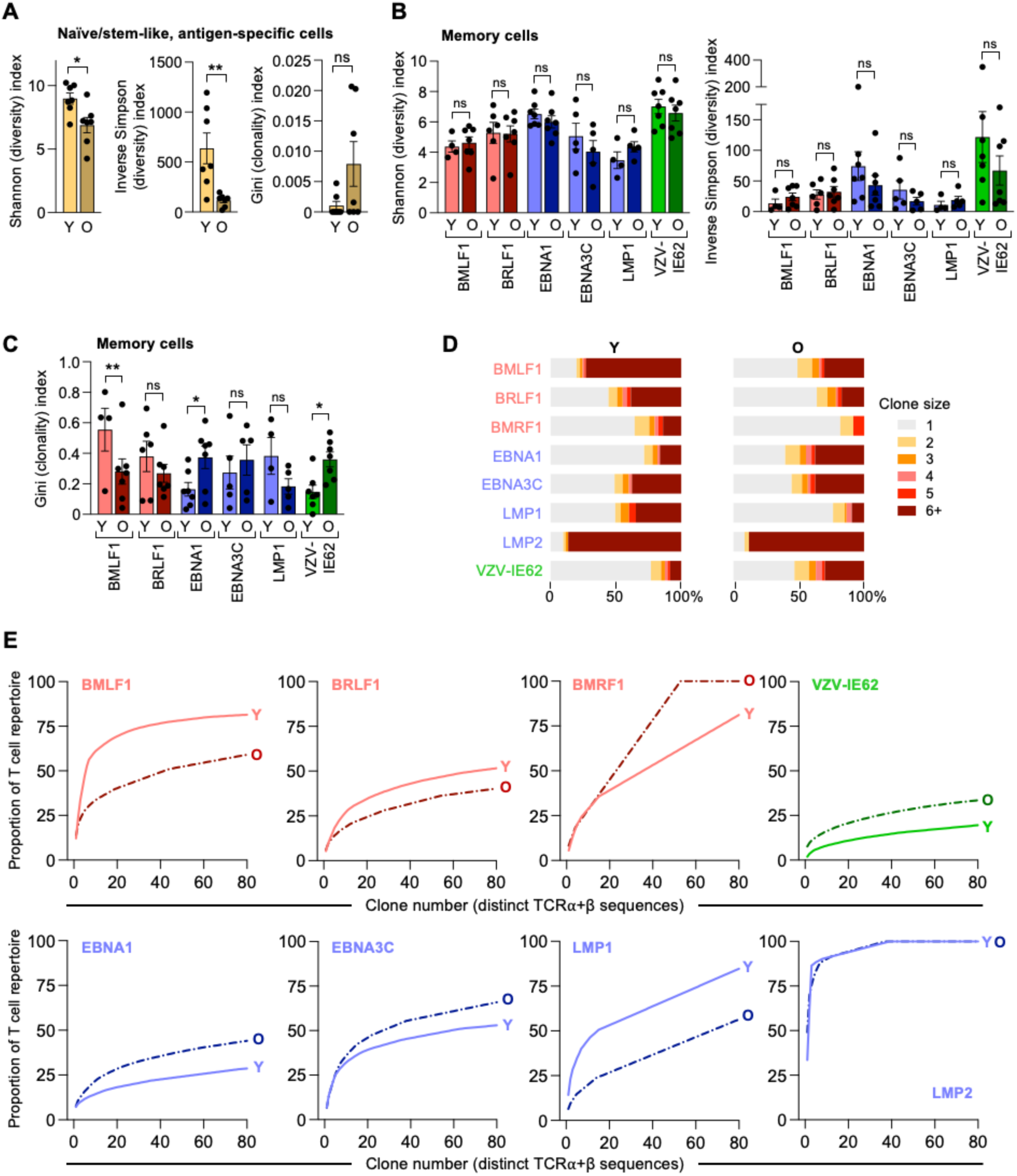
Memory T cell receptor (TCR) diversity of antigen-specific T cells is maintained with aging. A. T cell receptor diversity and clonality are shown as Shannon (left), Inverse Simpson (middle) and Gini indices (right) in antigen-specific CD8^+^ T cells mapping to the naïve/stem-like cluster (see Fig. 5A). Only cells with TCRα and TCRβ sequences in single cell-sequencing data were used for the calculation. Due to low cell numbers, indices were calculated on the combined populations of all antigen-specific T cells with naïve/stem-like phenotype. **B**. T cell diversity indices as in (A) but calculated on antigen-specific T cells with a memory phenotype comparing cells from young (Y) and older (O) individuals. **C**. Gini index as in (A) to estimate T cell clonality in antigen-specific T cells with a memory phenotype. **D**. Clonal size distributions for each antigen specificity are shown as stacked bars for Y and O adults. **E**. Numbers of distinct antigen-reactive TCR chains ordered by decreasing clonal sizes plotted against the cumulative space they occupy. The numbers of distinct antigen-reactive TCR chains from Y adults as shown in Fig. 4G is compared to results from O adults (dotted lines). Data show the mean ± SEM (A-C). All datapoints represent distinct biological replicates. O datapoints were contrasted to Y data described in Fig. 4 and Supplementary Fig. 6. Data were compared by two-tailed, unpaired *t*-tests (A) or two-way ANOVA with Šídák’s multiple comparisons test (B, C). *P<0.05, **P<0.01. ns, not significant.

## DISCUSSION

Here, we describe that CD8^+^ T cells specific for different EBV-derived peptides differ in the changes that they undergo with aging. These differences correlate with differentiation states that T cells have developed with the initial infection and that are preserved into young adulthood. In general, T cells that are specific for lytic EBV antigens are already more biased towards effector end-differentiation than those specific for latent antigens in young adults. Conversely, T cells specific for latent antigens undergo more progressive changes with age as determined by phenotypic analyses, expression of transcription factors, and transcriptomic signatures. At a more granular level, heterogeneity of age-associated changes in EBV-specific T cells exists even within the sets of lytic and latent antigens that we tested. While a common trajectory was a loss in the expression of genes related to adaptive immunity and a gain in innate, NK cell-related immunity, age-associated changes in the transcriptome were not widely shared between T cells specific for peptides derived from different EBV proteins, and each antigen-specificity had its own dominant aging signature.

How immune memory is maintained over lifetime is central to healthy aging and has important implications for vaccine strategies. While significant progress has been made in the last decade in describing age-associated changes in the global T memory cell compartment, one of the major limitations of current datasets is that they do not allow conclusions at the level of antigen-specific T cells. Studies at the population level of CD8^+^ T cells have described shifts to more differentiated cells, i.e., effector, TEMRA, and T cells expressing activation or exhaustion markers^38^. While phenotypic studies alone cannot completely capture the heterogeneity of CD8^+^ memory T cells^3^, omics approaches including single cell transcriptome studies have significantly expanded our insights into human immune aging in recent years. Datasets on PBMCs of >100 healthy individuals have identified an age-associated CD8^+^ T cell subpopulation that was characterized by the expression of granzyme K^39,40^. A CD8^+^ memory T cell populations marked by NKG2C^+^ GZMB^−^, however, decreased with age^41^. While single cell 5’-RNA-seq allows to identify TCR sequences and therefore assign transcriptome signatures to clonal specificities, conclusions from cross-sectional studies on the durability of antigen-specific T cells are difficult given the immense TCR diversity of even antigen-specific T cells across individuals. Longitudinal studies are usually limited to up to 2 years after vaccination^24,42^. The currently longest longitudinal study by Sun *et al.*^43^ only focused on changes in the TCR repertoire over a timespan of ∼9 years using the Baltimore Longitudinal Study of Aging.

Elegant, murine studies provided surprising insights into how antigen-specific memory T cells evolve with repeated restimulation over lifetime. Soeren *et al.*^7^ used iterative stimulations and adoptive transfers in mice and showed that mouse memory T cells remained fully functional over 51 immunizations without evidence of senescence or exhaustion, across a time span of >10 years, far exceeding a mouse’s lifespan. Functionality was sustained despite chromatin remodeling at exhaustion markers but was dependent on sufficient rest between stimulation events. Analogous to this mouse study, van de Sandt *et al.*^44^ examined how epitope-specific T cells evolve across the human lifespan by examining the phenotype and transcriptome of CD8^+^ T cells directed at the prominent influenza matrix epitope HLA-A*02:01-M1_58–66_ over lifetime. Epitope-specific CD8^+^ T cells and total CD8^+^ T cells displayed different phenotype profiles with TEMRA cells increasing in the total CD8^+^ T cell population with age but not in influenza-specific T cells suggesting that influenza M1 stimulation does not trigger terminal differentiation per se. The transcriptome of older influenza-specific T cells correlated with less-differentiated cell states, lacking gain in effector function and expression of cytotoxic molecules, very much in contrast to total CD8^+^ T cells. Of note, there was no evidence for the expression of cellular senescence or exhaustion genes in older adults although cells lost activation and proliferative capacity. These age-associated changes in transcriptomes were correlated with a shift in the TCR repertoire suggesting that they were not due to T cell aging but recruitment of a new set of naïve T cells into the memory pool.

To probe the evolution of human memory T cells over lifetime and determine whether they develop genetic hallmarks of cellular senescence or exhaustion, we elected to examine several epitope-specific CD8^+^ T cells in EBV infection. Notable differences to the influenza matrix epitope system are that EBV is a chronic latent infection while the re-exposure history to the matrix protein is more difficult to standardize. The matrix protein is largely absent in component vaccines and re-exposure therefore depends on infections that can be clinically silent. Moreover, the EBV system allows to compare the age-dependent evolution of CD8^+^ T cells specific for multiple lytic and latent antigens that were primed simultaneously early in life. The primary immune response in acute mononucleosis constitutes a dramatic expansion of CD8^+^ T cells specific to lytic proteins that can make up 50% of all CD8^+^ T cells.^9^ Cells responding to latent epitopes are present at lower frequencies contributing up to 5% of CD8^+^ T cells. After the acute infection, most effector T cells are culled and memory T cells specific to lytic and latent antigens develop at a broadly similar immunodominance hierarchy to those seen in effector T cells. Importantly, Hislop *et al.*^12^ found that by the time the primary EBV infection resolves, CD8^+^ memory T cells have developed distinct differentiation states dependent on their epitope specificity. On average, responses to lytic antigens have a more progressed effector phenotype. In our study population who were in their early adulthood up to 40 years old, we observed that EBV-specific T cells still differed in their differentiation states based on the epitope of the protein that they recognized. These states loosely correlated with the classification into lytic and latent proteins, however, transcriptome and chromatin structure data yielded a more complex picture with each antigen specificity having a unique signature. Given the data by Hislop *et al*. that these differences in T cell differentiation states are established early during infection, they are obviously maintained over more than a decade and gave us the opportunity to examine whether they have different aging trajectories, e.g., whether more frequent and more differentiated T cells are more susceptible to develop features of exhaustion or cellular senescence. This was not the case, and transcriptome changes were more pronounced for less differentiated specificities, including an increased expression of granzyme B, in contrast to recently reported data on cells specific for the influenza matrix protein^44^. Granzyme K, recently described as a common signature of aging^40,41,45^, was widely expressed in our EBV-specific cells, irrespective of age.

The memory T cell compartment is highly dynamic. Individual cells have limited lifespans, certainly considerably shorter than the duration of immunological memory. Memory persistence is therefore not conferred by the longevity of individual memory cells but by a population of cells that are individually short-lived^46^. This scenario should favor a selection for fitness in the repertoire of EBV-specific T cells that could be different for different epitopes. Indeed, initial observations by Rickinson and Hislop suggested that effector T cells specific for lytic antigens contract more than those specific for latent antigens in the early stages after infection^12^. We observed that frequencies of all EBV antigen specificities are maintained into older age. TCR diversity was variable between different specificities with BMRF1- and LMP1-reactive T cells having only few clonally expanded memory cells. Previously reported longitudinal studies up to 5 years have reported persistence of individual clonotypes^47–49^. In our cross-sectional comparison, we see a lack of singletons expressing a naïve/stem-like phenotype with age, in line with a general loss of bulk naïve CD8^+^ T cells^50^. Either these singletons are truly lost or they are expanded and recruited into the memory compartment – a phenomenon that has been reported after infectious mononucleosis in young adults^51^ and for VZV-specific CD4^+^ T cells after vaccination in older adults^52^. Indeed, if we restrict our analysis to only phenotypically defined memory T cells, we observe less clonality and more diversification of the TCR repertoire for some EBV-epitope specific T cells specificities with age.

Our data support the notion that controlling for T cell heterogeneity including antigen specificity is needed to define age-associated changes that are clinically relevant. Understanding whether and how age affects diversity, function, and durability of memory T cells has implications for designing effective vaccination strategies in older adults, who are exceedingly more vulnerable to infections^1,2^, as well as for optimizing T cell therapies in cancer patients. Specifically, understanding fidelity of EBV-specific T cells is relevant as EBV is a major oncogenic stimulus for malignancies, many of which are more often diagnosed in older adults^18^. In EBV-related nasopharyngeal carcinoma or Hodgkin’s lymphoma, adoptive transfer of EBV-specific T cells is successfully pursued as cellular immunotherapy^53–55^. In this approach, autologous or allogeneic T cells are stimulated *ex vivo* with EBV antigens LMP1, LMP2, and/or EBNA1, expanded, and infused into the cancer patient. Total EBV-specific T cell numbers and their phenotype show some association with stable disease^56^ and extended overall survival^54^, but further optimization of the transferred EBV-specific T cells will be beneficial to reduce variability and improve efficacy for this cell therapy approach.

## METHODS

### Human subject study population

Peripheral blood mononuclear cells (PBMCs) from leukoreduction system chambers of 38 blood or platelet donors were purchased from the Mayo Clinic Blood Donor Center, Department of Laboratory Medicine and Pathology - Component Laboratory. Samples were de-identified except for age and sex information. Samples from male and female individuals were used. Young adults were 21 to 40 years old, older adults were 65 years or older. In addition, PBMCs from 27 healthy volunteers without acute or active chronic disease, and no history of cancer or autoimmune disease were recruited from a healthy aging registry at Stanford University. Chronic diseases were permitted if controlled by medication. Subject information on samples used in key experiments is summarized in Supplementary Table 1. Studies involving human subjects were approved by the Stanford University and Mayo Clinic Institutional Review Boards.

### Primary human cells

PBMCs were purified by density centrifugation with Lymphoprep (StemCell Technologies, #07861). Alternatively, T cells were isolated with EasySep Human T Cell Enrichment Kit (StemCell Technologies, #19051). Cells were either directly used or frozen in aliquots in Bambanker freezing medium (Lymphotec Wako Chemicals, #302-14681). For ATAC-seq, fresh cells were used; for flow cytometric phenotyping analyses, fresh or frozen cells; for single cell sequencing, frozen cells that were rested before experiments. Total CD8^+^ T cells were isolated using EasySep Human CD8^+^ T Cell Isolation Kit (StemCell Technologies, #17953). Screening for HLA-A*02-positive individuals was performed with an anti-human HLA-A2 Antibody (BioLegend, # 343326). For single cell sequencing, total CD8^+^ T cells were purified as above, resuspended in RPMI-1640 medium (Sigma Aldrich, #R8758) supplemented with 5% filtered human AB serum (Sigma Aldrich, #H4522), 100 U/mL penicillin, and 100 U/mL streptomycin (Sigma Aldrich, #P0781) and rested in a 37°C humidified incubator (5% CO_2_, atmospheric O_2_) for 16 hours. Cells were collected and washed before subjecting them to BEAM-T and antibody staining.

### Flow cytometry

For phenotyping of EBV antigen-specific CD8^+^ T cells, purified CD8^+^ T cells were first stained with tetramers against EBV antigens (BMLF1, BRLF1, LMP2, EBNA3C, see Supplementary Table 2). About 5 million cells were stained with 5 uL of each tetramer (containing 0.2 ug MHC monomer conjugated to 0.05 ug streptavidin and fluorophore) and PBS containing 2% fetal bovine serum (FBS) (GeminiBio, #900-108) in a total volume of 50 uL. After 45 minutes of staining at 4°C, diluted antibodies against cell surface proteins (see Supplementary Table 2) and a viability dye (LIVE/DEAD Fixable Blue, Thermo Fisher Scientific, #L23105) were added. Cells were stained for an additional 30 minutes at 4°C. For the cell surface panel, cells were washed with PBS containing 2% FBS and fixed with Cytofix fixation buffer (BD Biosciences, #554655) for 30 minutes at 4°C, washed and subjected to spectral flow cytometry. For the intracellular transcription factor panel, cells were washed with PBS containing 2% FBS and fixed with the eBioscience Foxp3 / Transcription Factor Staining Buffer Set (Thermo Fisher Scientific, #00-5523-00) according to the manufacturer’s instruction. Briefly, cells were fixed for 30 minutes at 4°C, washed and permeabilized. Antibodies against intracellular proteins (see Supplementary Table 2) were diluted in permeabilization buffer and staining was performed for 45 minutes at room temperature in the dark. After washing, cells were subjected to spectral flow cytometry. A 5-laser Cytek Aurora (Cytek Biosciences) running SpectroFlo v3 was used to collect data. UltraComp eBeads compensation beads (Thermo Fisher Scientific, #01-2222-42) were used for single staining controls. At least 1 million cells were collected, and at least 100 tetramer-positive cells were used for analyses. A representative gating strategy is shown in Fig. 1C.

### Flow cytometry analyses

FlowJo v10 was used to analyze data. Live, single CD3^+^ CD8^+^ T cells were analyzed for tetramer-staining or classical phenotypic subset markers: naïve (CD45RA^+^ CCR7^+^ CD28^+^), central memory (CD45RA^−^ CCR7^+^ CD28^+^), effector memory (CD45RA^−^ CCR7^−^) and TEMRA cells (CD45RA^+^ CCR7^−^ CD28^−^). Proteins of interest were analyzed in tetramer-specific cells and each subset. Representative gating schemes are shown in Supplementary Fig. 1C-E.

For clustering and reference mapping analyses, samples were randomly down-sampled to 50 000 CD8^+^ T cells per individual by the FlowJo plugin DownSampleV3. Channel values of these cells and all tetramer-positive cells were exported from FlowJo and imported into Seurat^57^ v4.2.1 for in-depth analyses. The raw counts were merged and transformed using centralized log ratio (CLR) across cells using the NormalizeData function. Data were scaled and principal component analysis (PCA) was performed using ScaleData and RunPCA function, respectively. Samples from different experiments were integrated by diagonalized canonical-correlation analysis (CCA) that identifies anchors to act as guides for dataset integration using the function FindIntegrationAnchors and IntegrateData^57^. The integrated matrix underwent scaling and PCA calculation generating the Uniform Manifold Approximation Projection (UMAP) along with a uniform manifold optimization (uwot) model for reference mapping via the RunUMAP function. Utilizing the PCA, the graph-based Louvain clustering was performed via FindNeighbours and FindClusters function from 0.2 to 1 to identify an optimal cluster resolution. Data visualization was performed via Seurat built-in functions and custom scripts.

To map tetramer-positive cells to the reference dataset, tetramer-positive samples were pre-processed as described above including CLR transformation, identification of variable features using NormalizeData and FindVariableFeatures. Subsequently, tetramer-positive cells were aligned with the reference to establish anchors, facilitating the transfer of cell type labels from reference to the samples using the FindTransferAnchors and TransferData function, respectively^57^. We summarized the overall distribution of tetramer-positive cells across reference clusters. These proportions were utilized to compute the principal component for proper representation of the samples in lower dimension.

### ATAC-seq and analysis

CD8^+^ tetramer-positive cells and CD8^+^ populations of central memory (CD45RA^−^ CCR7^+^), effector memory (CD45RA^−^ CCR7^−^) and TEMRA cells (CD45RA^+^ CD28^−^) were collected via FACS. Representative gating is shown in Supplementary Fig. 3B. ATAC-seq was performed as previously described^24^. We aligned ATAC-seq reads to the hg38 reference genome and normalized data as described previously^22,23^. Peaks that were present in at least 6 samples were included. Peaks with low read counts across sample groups were removed using the filterByExpr function^58^. PCA was calculated using regularized log transformed counts from the top 5000 variable peaks using plotPCA function in DESeq2. Peak counts were normalized using conditional quantile normalization (CQN) taking into consideration GC bias and length of the peaks for each sample. The normalized data were modelled for mean-variance trend for each gene incorporating the sample weight adjustment using voomWithQualityWeights function in limma^59^. Differential peaks (adjusted *P*-value <0.05) were calculated across all populations by fitting each contrast to the linear model and employing empirical Bayes moderation. Differential peaks were clustered using K-means clustering with the default parameters. Clustered differential peaks were used for TF binding site (TFBS) prediction using HOMER^60^. TFBS prediction was also performed on all peaks using chromVAR package^25^. Peaks were annotated using ChIPseeker with region range of 1 kb across the transcription start site (TSS) and using TxDb.Hsapiens.UCSC.hg38.knownGene annotation^61^. Genes that were annotated as promoter or intragenic peaks were selected for enrichment analyses using EnrichR^62^ focusing on Gene Ontologies, Reactome, BioPlanet, KEGG and ChEA gene sets. The averaged bigwig files were used to visualize peak profiles using Integrated Genome Viewer (IGV)^63^.

### Barcode Enabled Antigen Mapping (BEAM)-T and single cell sequencing

MHC monomers (Chromium Human MHC Class I A*02:01 Monomer Kit, 10x genomics, #1000542) were loaded with EBV peptides (GenScript) and incubated with the PE-containing BEAM conjugate (Chromium Single Cell 5’ BEAM Core Kit PE, 10x genomics, #1000539) according to the manufacturer’s instructions. Peptides for BMLF1 (GLCTLVAML), BRLF1 (YVLDHLIVV), BMRF1 (TLDYKPLSV), BALF4 (FLDKGTYTL), EBNA1 (FMVFLQTHI), EBNA3C (LLDFVRFMGV), LMP1 (YLQQNWWTL) LMP2 (CLGGLLTMV), VZV-IE62 (ALWALPHAA), and VZV-IE63 (RLVEDINRV) were custom synthesized by GenScript. Loading of each peptide onto the MHC monomer was confirmed prior to use. A negative control peptide was provided within the kit and included in all analyses.

Magnetic-activated cell sorting (MACS)-purified CD8^+^ T cells from peripheral blood were subjected to BEAM-T staining according to the manufacturer’s instructions. Four to five individuals per experiment (3 experiments total) were processed. Briefly, about 3-4 million CD8^+^ T cells were blocked with Fc receptor blocking solution (Human TruStain FcX, BioLegend, #422302) for 10 minutes on ice and stained with BEAM-T assemblies for 15 minutes on ice, followed by cell surface marker antibodies plus unique hash-tagged CITE antibodies for 30 minutes on ice. Cells were washed 3 times with PBS containing 2% FBS before 7-AAD viability staining solution (BioLegend, #420404) was added and cells were subjected to FACS (BD FACSAria 4-laser digital flow cytometer with FACSDiva v8 software). Cell sorting was performed by the Mayo Clinic Microscopy and Cell Analysis Core Flow Cytometry Lab. Live, single CD3^+^ CD8^+^ PE^+^ cells representing BEAM-T^+^ antigen-specific cells and PE^−^ cells representing total CD8^+^ T cells were sorted. After FACS, cells were counted and equal cell numbers of PE^+^ and PE^−^ cells from each individual were pooled. The pooled sample was then stained with diluted TotalSeq-C cell surface marker antibodies (BioLegend, Supplementary Table 2) in PBS containing 2% FBS for 30 minutes at 4°C. Cells were washed twice with PBS containing 2% FBS, and cell viability was determined with Trypan Blue. Cell viability was consistently >90%. Cells were subjected to high-throughput single cell capture via the Chromium X controller (10x genomics), Chromium Next GEM Single Cell 5’ HT Kit v2 (Dual Index, 10x genomics, #PN-1000377) and gel beads (10x genomics, #1000376). About 40,000 cells were targeted for cell capture for each side of the HT chip. Four libraries were generated from each captured sample: 5’ gene expression, feature barcodes (for TotalSeq-C antibodies including hashtag antibodies and cell surface marker, 5’ Feature Barcode Kit, 10x genomics, #PN-1000541), TCR sequences (Chromium Single Cell V(D)J Amplification kit, 10x genomics, #PN-1000252) and BEAM-T antigen specificity. All captured samples and libraries were sequenced in one flow cell using a 100 cycle kit (paired-end) on a NovaSeq 6000 S4 (Illumina) targeting 20,000 read pairs/cell for gene expression, 5,000 read pairs/cell for feature barcodes, 5,000 read pairs/cell for TCR sequences, and 5,000 read pairs/cell for BEAM-T. Cell capture and library construction was performed by the Mayo Clinic Medical Genome Facility - Genome Analysis Core and sequencing was performed by the San Diego Nathan Shock Center Heterogeneity of Aging Core.

### Single cell sequence analyses

Quality assessment of the fastq reads were conducted using FastQC. Subsequently the four modalities (gene expression, GEX; CITE antibodies, ADT; T cell receptor sequences (VDJ), and antigen specificity by BEAM-T, Ag) were processed using 10x Genomics Cell Ranger v7.1.0 multi for each experiment^64^. GEX was aligned to the Cell Ranger-provided GRCh38-2020-A reference, and VDJ reads were aligned to Cell Ranger-provided vdj-GRCh38-alts-ensembl-7.1.0 reference. For ADT and Ag, unique feature barcodes were utilized to extract the specific unique molecular identifier (UMI) associated with both modalities. Sample demultiplexing was carried using hash-tagged CITE antibodies using 10x Genomics cellranger v.7.1.0 multi^64^. We noticed that about half of the good quality cells that were identified in CellRanger could not be demultiplexed via antibody-based demultiplexing due to the low UMIs of hash-tagged CITE antibodies. To overcome this challenge, variant-based demultiplexing based on GEX was implemented for each sample using souporcell^65^. Comparison of the hash-tag-based demultiplexing and variant-based demultiplexing revealed very high accuracy (>0.98), sensitivity (>0.97) and specificity (>0.99) across all experiments.

Poor-quality cells were excluded based on the following criteria: cells with <200 or >5000 expressed genes, >10% UMIs mapping to mitochondria-encoded transcripts and >10,000 ADT UMIs. Additionally, DoubletFinder was used to remove cell doublets. To differentiate antigen-specific cells from the bulk CD8^+^ T cell population, Ag UMIs were modeled on beta distribution by 10x Genomics Cell Ranger v7.1.0 multi^64^ to calculate the probability of antigen specificity (known as antigen specificity score) in comparison to the negative control peptide provided by 10x genomics. Cells with antigen specificity scores of >50 were classified as putative antigen-specific CD8^+^ T cells. Cells with scores of<50 and with low Ag (BEAM-T) UMIs (<40 UMIs for experimental batch 1 and <75 UMIs for experimental batches 2 and 3) were classified as bulk CD8^+^ T cells. To annotate the identity of antigen for each cell, the antigen with the highest antigen specificity score was selected. To account for possible cross-reactivity, also antigens that fall within an antigen specificity score range of 10 from the maximal antigen specificity score were selected. Cells specific to BALF4 and VZV-IE63 were too infrequent and, thus, these specificities were excluded from the analysis. The bulk T cell population and antigen-specific samples were processed similarly for GEX and ADT. GEX samples were individually normalized using SCTransform v2^66^, while CLR transformation was applied for ADT counts. Subsequently, samples from different experimental batches were integrated using diagonalized CCA to identify mutual nearest neighbor that acted as anchors for integration employing the functions FindTransferAnchors and IntegrateData. Integrated data from GEX and ADT were scaled and PCA was calculated to integrate the modalities using weighted nearest neighbors (WNN) through FindMultiModalNeighbors function. Based on the WNN, we generated the combined UMAP using the RunUMAP function. Further clustering was performed using smart local moving (SLM) algorithm^57^. For data visualization, we addressed dropout in GEX and ADT via Markov affinity-based graph imputation of cell (MAGIC) that denoises the matrix and imputes values across similar cells via data diffusion^67^. Cell types were annotated for each cluster based on ADT cell surface markers and GEX gene markers identified using Seurat built-in FindAllMarkers function. Visualization was performed via Seurat built-in functions and custom scripts^57^.

### Pseudobulk differential expression and K-means clustering

Raw counts of antigen-specific samples were aggregated by summing the UMIs from their respective single cells utilizing Seurat’s built-in function AggregateExpression^57^. Antigen-specific samples with fewer than 20 cells were not considered. Additionally low-expressing genes with counts fewer than 10 within the group were filtered out using the filterByExpr function^58^. Raw counts were then regularized log (rlog) transformed for computing principal components, followed by batch effect removal using plotPCA and removeBatchEffect function, respectively^58,59^. Differentially expressed genes (DEGs) for each antigen-specific T cell comparison between older and young individuals as well as between antigen-specific T cells of young individuals were calculated. To identify the DEGs, the mean-variance trend was modelled on the log-transformed counts per million (logCPM) value of each gene, incorporating the sample weight adjustment via voomWithQualityWeights in limma^59^. DEGs were calculated by fitting contrasts to the model and employing empirical Bayes moderation with log_2_ fold change +/– 0.5 and adjusted *P*-value <0.05. DEGs were clustered using the k-means function with default parameters. Enrichment analyses were performed using EnrichR^62^ focusing on Gene Ontologies, Reactome, BioPlanet, KEGG and ChEA gene sets. Gene set enrichment analysis (GSEA) for adaptiveness and innateness genes^36^, exhaustion-associated genes (GSE9650, GSE41867) and cellular senescence-associated genes (GO:0090398, Reactome R-HSA-2559583, SenSig^68^, SenMayo^69^) was performed by FGSEA v1.20.0^70^. UpSet plots were generated with Intervene^71^ and heatmaps with Morpheus (https://software.broadinstitute.org/morpheus).

### High-dimensional tetramer-associated T cell receptor sequence analysis

We downloaded publicly available high-dimensional tetramer-associated T cell receptor sequence (TetTCR-SeqHD)^21^. TetTCR-SeqHD data include targeted GEX, ADT, TCR and antigen-specificity against 282 antigens. We extracted EBV-specific CD8^+^ T cells with specificities against BMLF1, BRLF1, BZLF1, LMP1, LMP2, LMP2A, EBNA3A. We processed data as described for self-collected samples elaborated on in the preceding section.

### T cell receptor diversity and clonality

To calculate TCR diversity and clonality, we extracted nucleotide sequences of complete TCR α and β chains (including CDR1, CDR2, and CDR3). We adopted the Inverse Simpson index and Shannon index to calculate TCR diversity, the Gini index was used to calculate TCR clonality using custom scripts. The Inverse Simpson index (D) was calculated as 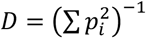 with *p* being the proportion of the i-th clones. The Shannon index (H) was calculated as *H* =∑(*p_i_*log_2_*p_i_*). The Gini index (G) was calculated as 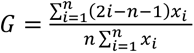 where i is the rank of values in ascending order of clones, n is the number of clones and x_i_ is a clone in the i-th rank.

### Statistical analyses

For statistical association of cluster proportions with antigen specificity and age, we characterized each sample as a probability vector, whose components represent the relative cell frequency in each of the clusters and the cardinality represents the number of distinct clusters. To examine the association between the probability vector and the antigen specificity, we employed linear regression analysis between the individual component of the probability vector, i.e., the relative cell frequency in a cluster, and antigen by treating the probability component as outcome and antigen specificity as independent variable of interest. We adjusted gender in the regression analysis. The association with the entire probability vector is summarized by the sum of squares of all z scores corresponding to the regression coefficient of the antigen specificity from the linear regression analyses and tested via permutation test permuting the antigen specificity. A similar method was used to assess the association of age within each antigen specificity with gender adjustment.

All other statistical analyses were performed via Prism software 10 (GraphPad). Data are presented as mean with error bars indicating standard error of the mean (SEM), unless otherwise indicated. Unpaired, two-tailed Student’s t-tests were used when comparing two groups. Two-way ANOVA with Šídák’s multiple comparisons test was used for multigroup comparisons. A P-value less than 0.05 was considered statistically significant. Significance levels (*P < 0.05, **P<0.01 and ***P<0.001) are indicated in figures.

## Data availability

ATAC-seq and single-cell RNA-seq data have been deposited at GEO (accession numbers not yet available) and are publicly available as of the date of publication. This study also analyzed existing, publicly available datasets (GEO GSE101609, dbGaP: phs002441.v1.p1).

## Code availability

This paper does not report original code. All employed code is open source and described in the respective method sections.

## Acknowledgements

We thank Elsa Molina and the San Diego Nathan Shock Center for their support on single cell-sequencing, and Katayoun (Kathy) Ayasoufi and Aaron Johnson for assistance with Cytek spectral flow cytometry. This work was supported by National Institutes of Health (NIH) R01AG045779, R01AI108891, R01AI129191, and U19AI057266 (all to J.J.G.); NIH R01AR042527, R01AI108906, R01HL117913, and R01HL142068 (all to C.M.W.); T32AG049672 (to I.S.), by a Glenn Foundation for Medical Research Postdoctoral Fellowship in Aging Research (to I.S.), by a Mayo Clinic Robert and Arlene Kogod Center on Aging Career Development Award (to I.S.), and a pilot grant from the San Diego Nathan Shock Center (to I.S.). This study was supported with resources and the use of facilities at the Palo Alto Veterans Administration Healthcare System, the Mayo Clinic Microscope and Cell Analysis Core, and the Mayo Clinic Medical Genome Facility – Genome Analysis Core. The content is solely the responsibility of the authors and does not necessarily represent the official views of the NIH.

## Author contributions statement

I.S., A.J., C.M.W., and J.J.G. conceptualized the study. I.S. performed experiments with assistance of W.C. and H.O.. A.J. performed bioinformatic analyses with assistance of R.R.J.. B.H. performed ATAC-seq. L.T. consulted on statistical analysis. I.S., A.J., and J.J.G. analyzed and interpreted data and wrote the manuscript. All authors edited the manuscript.

## Materials & Correspondence

Supplementary information contains 7 supplementary figures and 2 supplementary tables. Correspondence and requests for materials should be addressed to Jörg. J. Goronzy.

