## Supplementary information for "Aging trajectories of memory CD8^+^ T cells differ by their antigen specificity"

### TABLE OF CONTENT

Supplementary Figures 1-7
Supplementary Tables 1-2

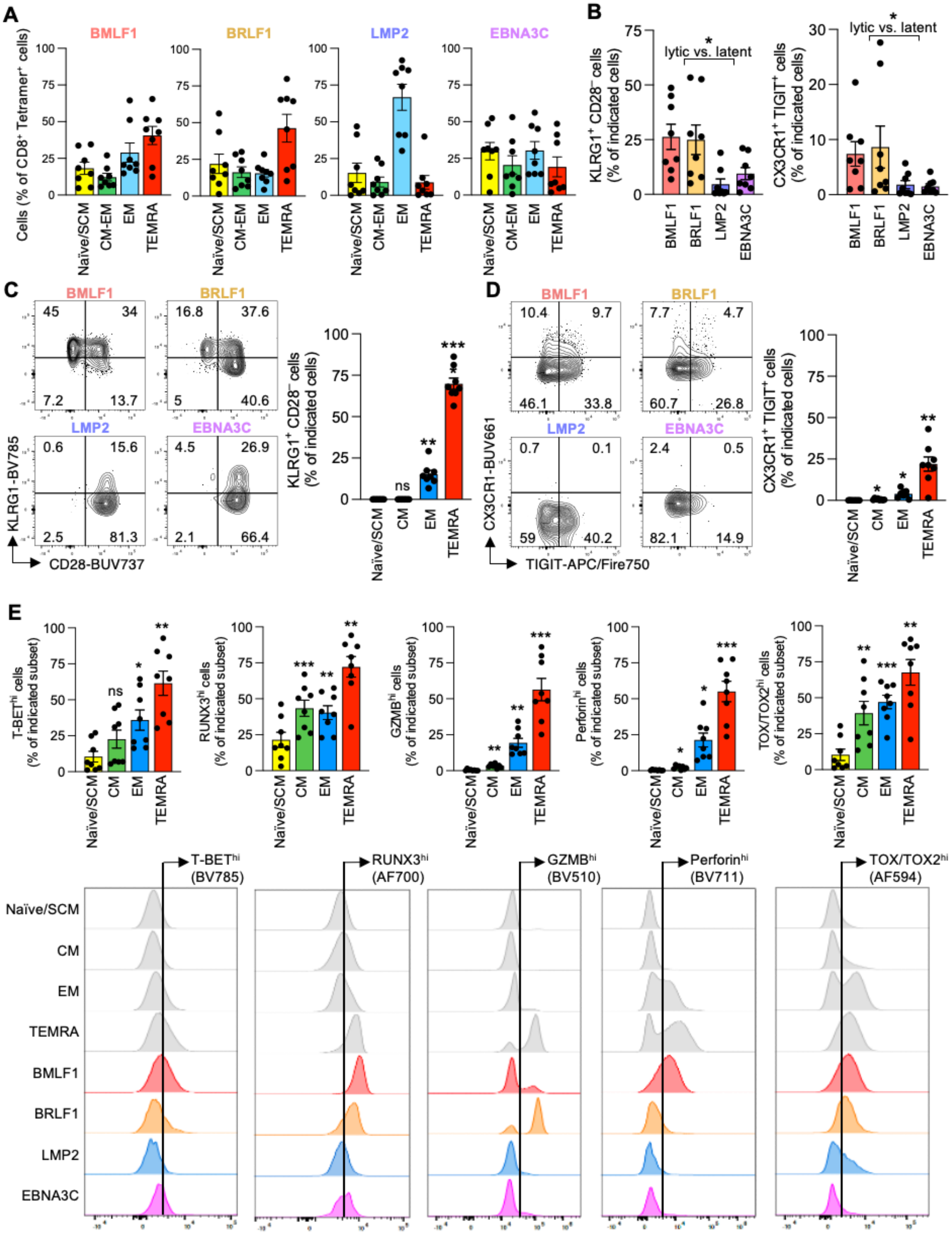

A
BMLF1
BRLF1
LMP2
EBNA3C
Cells (% of CD8+ Tetramer+ cells)
Naïve/SCM
CM-EM
EM
TEMRA
B
KLRG1+ CD28− cells (% of indicated cells)
CX3CR1+ TIGIT+ cells (% of indicated cells)
lytic vs. latent
C
KLRG1-BV785
CD28-BUV737
KLRG1+ CD28− cells (% of indicated cells)
D
CX3CR1-BUV661
TIGIT-APC/Fire750
CX3CR1+ TIGIT+ cells (% of indicated cells)
E
T-BEThi cells (% of indicated subset)
RUNX3hi cells (% of indicated subset)
GZMBhi cells (% of indicated subset)
Perforinhi cells (% of indicated subset)
TOX/TOX2hi cells (% of indicated subset)
T-BEThi (BV785)
RUNX3hi (AF700)
GZMBhi (BV510)
Perforinhi (BV711)
TOX/TOX2hi (AF594)
Naïve/SCM
CM
EM
TEMRA
BMLF1
BRLF1
LMP2
EBNA3C

**Supplementary Figure 1 (related to Figure 1): CD8$^{+}$ T cells against lytic EBV antigens express higher levels of end-differentiation markers**. **A**. Quantitation of EBV-specific cells across classical T cell subsets from reference mapping analyses in Fig. 1D. SCM, stem-like memory; CM, central memory; EM, effector memory; TEMRA, terminal effector memory with CD45RA expression. **B-D**. Flow cytometry quantification of indicated cell surface marker combinations indicative of advanced differentiation in antigen-specific CD8$^{+}$ T cells or classical T cell subsets. Representative flow cytometry plots are shown. **E**. Flow cytometry quantification of intracellular markers including transcription factors and effector molecules as in (B-D). Data show the mean ± SEM (A-E). All datapoints represent distinct biological replicates. Data were compared by two-tailed, unpaired *t*-tests (B) or one-way ANOVA with Šídák's multiple comparisons test (C-E). *P<0.05, **P<0.01, ***P<0.001. ns, not significant.

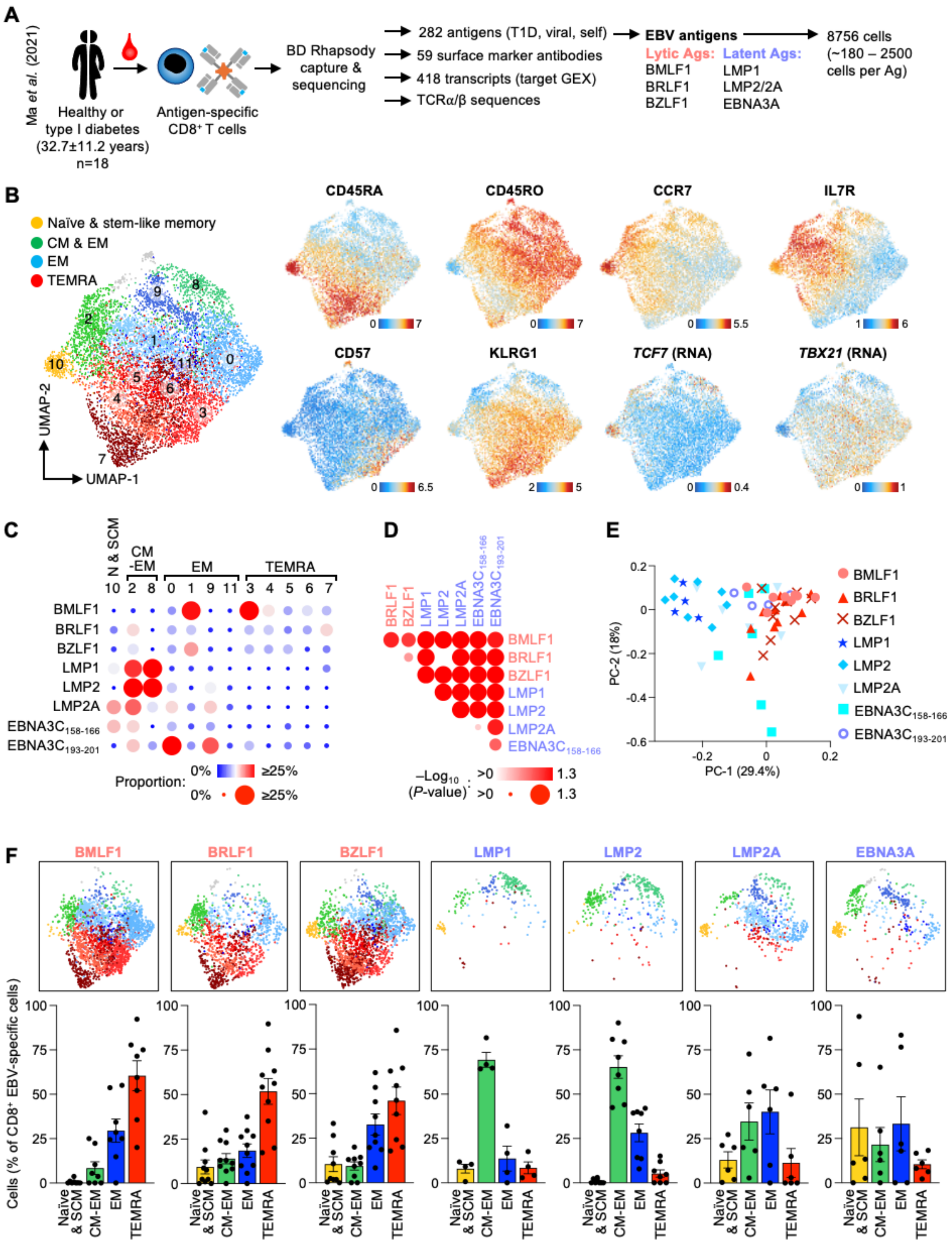
A
Ma et al. (2021)
Healthy or type I diabetes (32.7±11.2 years) n=18
Antigen-specific CD8⁺ T cells
BD Rhapsody capture & sequencing
282 antigens (T1D, viral, self)
59 surface marker antibodies
418 transcripts (target GEX)
TCRα/β sequences
EBV antigens
Lytic Ags:
BMLF1
BRLF1
BZLF1
Latent Ags:
LMP1
LMP2/2A
EBNA3A
8756 cells (~180 – 2500 cells per Ag)
B
Naïve & stem-like memory
CM & EM
EM
TEMRA
UMAP-1
UMAP-2
CD45RA
CD45RO
CCR7
IL7R
CD57
KLRG1
TCF7 (RNA)
TBX21 (RNA)
C
N & SCM
CM -EM
EM
TEMRA
BMLF1
BRLF1
BZLF1
LMP1
LMP2
LMP2A
EBNA3C158-166
EBNA3C193-201
Proportion: 0% ≥25%
D
–Log10 (P-value): >0 1.3
E
PC-1 (29.4%)
PC-2 (18%)
F
BMLF1
BRLF1
BZLF1
LMP1
LMP2
LMP2A
EBNA3A
Cells (% of CD8⁺ EBV-specific cells)
Naïve & SCM
CM-EM
EM
TEMRA

**Supplementary Figure 2 (related to Figure 1): EBV-specific CD8$^+$ T cell diverge phenotypically with large differences between T cells specific to antigens expressed in the lytic versus latent EBV life cycle**. **A**. Schematic of approach and data analyses of publicly available, single-cell sequencing data (dbGaP: phs002441.v1.p1) by Ma *et al*.[21] **B**. UMAP and KNN clustering of EBV-specific CD8$^+$ T cell sequencing data (left); selected marker expression to distinguish various T cell subsets and states (right). **C**. Bubble plot of EBV-specific T cell frequencies across clusters defined in (B). **D**. Probability vector analyses based on the distributions of antigen-specific cells across 11 subsets are compared. Significance levels are shown as bubble size and color gradient. **E**. PCA plot based on the subset distribution frequencies of antigen-specific T cell samples. **F**. Visualization of each antigen specificity as feature plot (top) and quantitation of EBV-specific cells across classical T cell subsets (bottom) in single-cell sequencing data as defined in (B). Data show the median (C) or mean ± SEM in (F). All datapoints represent distinct biological replicates.

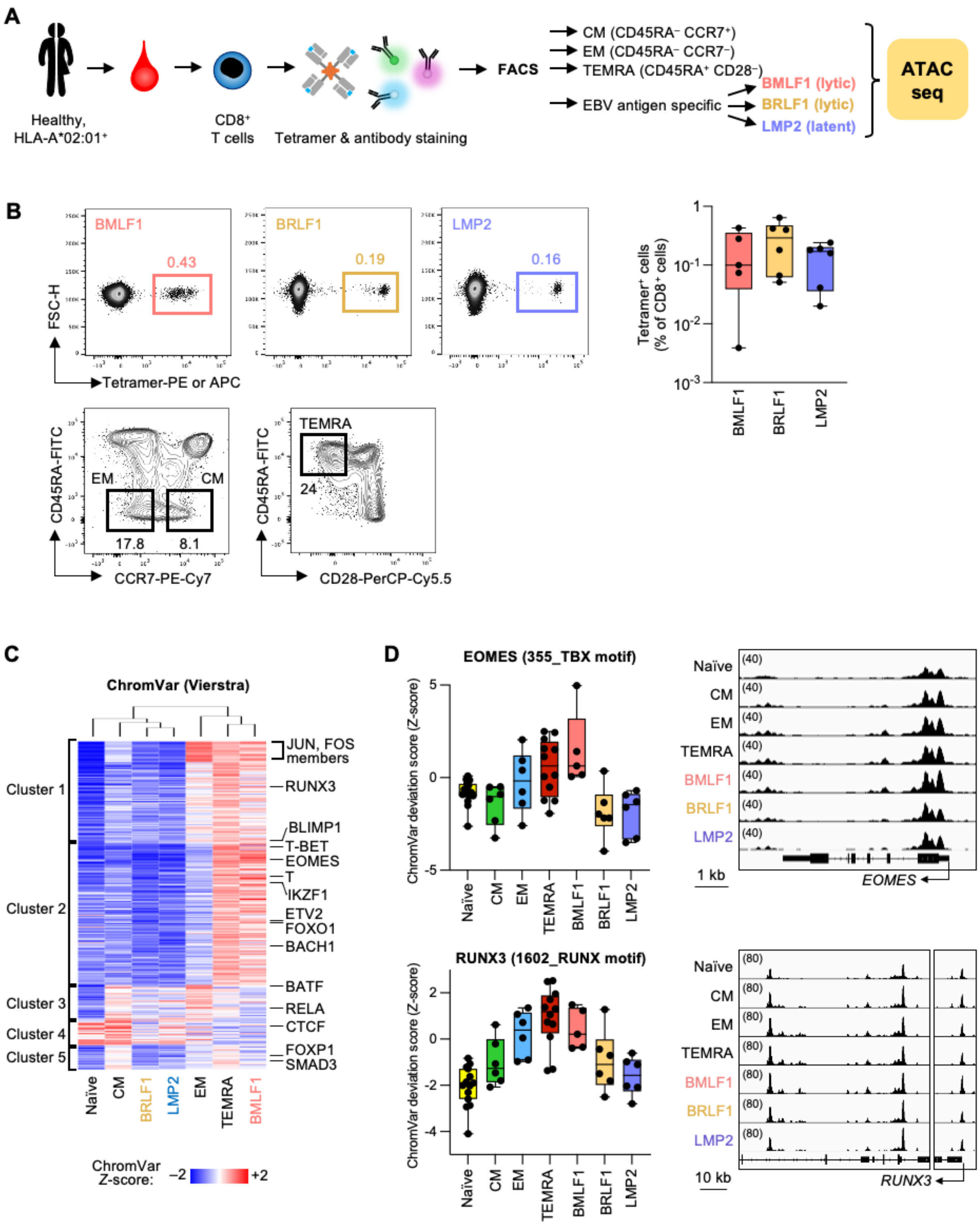
A
Healthy, HLA-A*02:01+
CD8+ T cells
Tetramer & antibody staining
FACS
CM (CD45RA− CCR7+)
EM (CD45RA− CCR7−)
TEMRA (CD45RA+ CD28−)
EBV antigen specific
BMLF1 (lytic)
BRLF1 (lytic)
LMP2 (latent)
ATAC seq
B
BMLF1
BRLF1
LMP2
0.43
0.19
0.16
FSC-H
Tetramer-PE or APC
Tetramer+ cells (% of CD8+ cells)
CD45RA-FITC
EM
CM
17.8
8.1
CCR7-PE-Cy7
TEMRA
24
CD28-PerCP-Cy5.5
C
ChromVar (Vierstra)
Cluster 1
Cluster 2
Cluster 3
Cluster 4
Cluster 5
JUN, FOS members
RUNX3
BLIMP1
T-BET
EOMES
T
IKZF1
ETV2
FOXO1
BACH1
BATF
RELA
CTCF
FOXP1
SMAD3
Naïve
CM
BRLF1
LMP2
EM
TEMRA
BMLF1
ChromVar Z-score: −2 +2
D
EOMES (355_TBX motif)
ChromVar deviation score (Z-score)
Naïve
CM
EM
TEMRA
BMLF1
BRLF1
LMP2
1 kb
EOMES
RUNX3 (1602_RUNX motif)
10 kb
RUNX3

**Supplementary Figure 3 (related to Figure 2): ATAC-seq signatures and corresponding transcription factor motif accessibilities distinguish the differentiation stages of antigen-specific T cells. A**. Schematic representation of experimental approach to collect EBV-specific T cells for ATAC-seq. **B**. Representative flow cytometry plots and gating strategy of EBV-specific T cells as well as bulk subsets (left). Quantitation of tetramer-positive T cells that were used for ATAC-seq (right). **C**. K-means clustering of ChromVar scores signifying inferred transcription factor (TF) motif accessibility in ATAC-seq peaks. **D**. ChromVar deviation scores for the EOMES and RUNX3 motifs (left) and ATAC-seq peak plots for these TF genes (right). Data show the median (B, D). All datapoints represent distinct biological replicates.

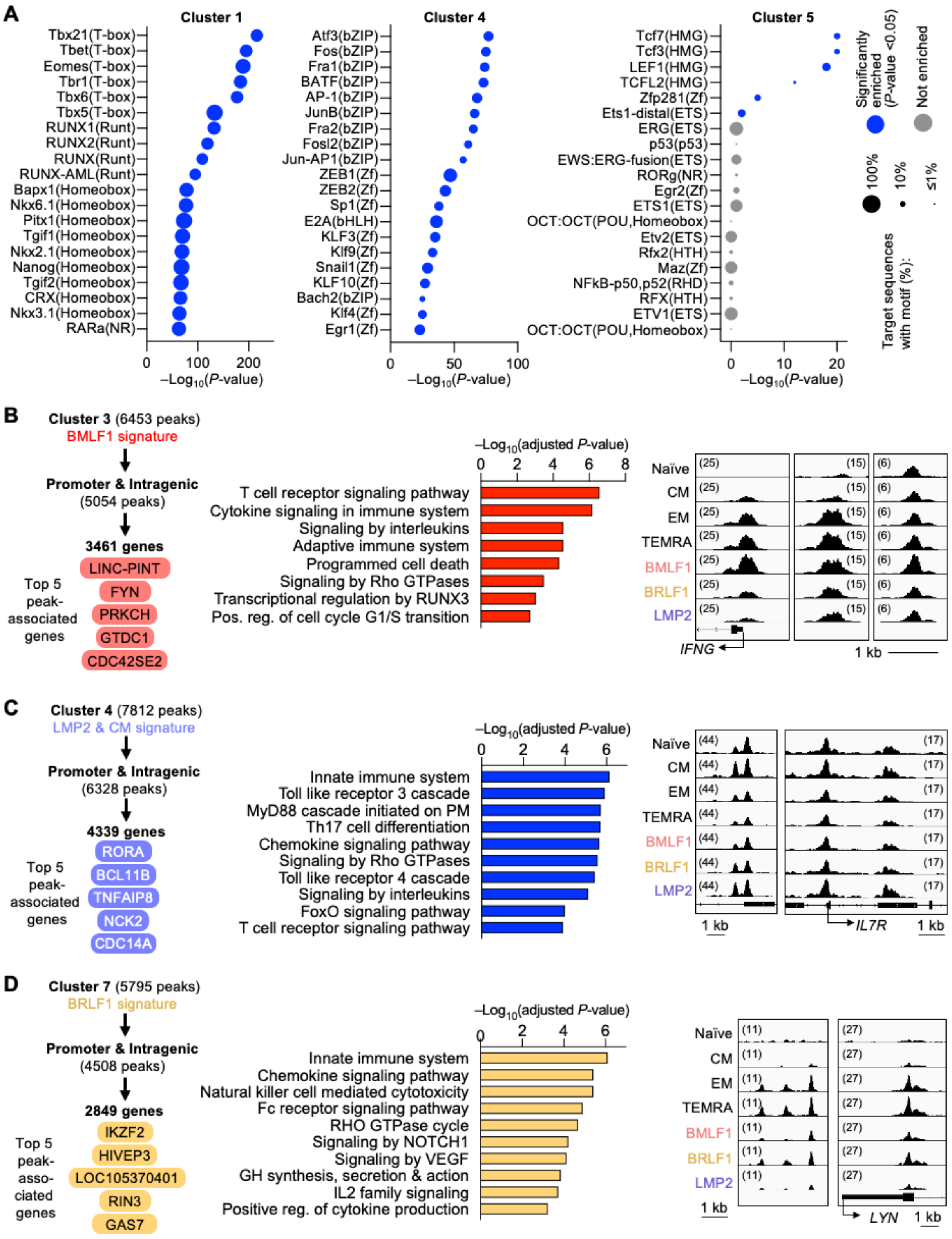

A
Cluster 1
Tbx21(T-box)
Tbet(T-box)
Eomes(T-box)
Tbr1(T-box)
Tbx6(T-box)
Tbx5(T-box)
RUNX1(Runt)
RUNX2(Runt)
RUNX(Runt)
RUNX-AML(Runt)
Bapx1(Homeobox)
Nkx6.1(Homeobox)
Pitx1(Homeobox)
Tgif1(Homeobox)
Nkx2.1(Homeobox)
Nanog(Homeobox)
Tgif2(Homeobox)
CRX(Homeobox)
Nkx3.1(Homeobox)
RARa(NR)
0 100 200
–Log10(P-value)
Cluster 4
Atf3(bZIP)
Fos(bZIP)
Fra1(bZIP)
BATF(bZIP)
AP-1(bZIP)
JunB(bZIP)
Fra2(bZIP)
Fosl2(bZIP)
Jun-AP1(bZIP)
ZEB1(Zf)
ZEB2(Zf)
Sp1(Zf)
E2A(bHLH)
KLF3(Zf)
Klf9(Zf)
Snail1(Zf)
KLF10(Zf)
Bach2(bZIP)
Klf4(Zf)
Egr1(Zf)
0 50 100
–Log10(P-value)
Cluster 5
Tcf7(HMG)
Tcf3(HMG)
LEF1(HMG)
TCFL2(HMG)
Zfp281(Zf)
Ets1-distal(ETS)
ERG(ETS)
p53(p53)
EWS:ERG-fusion(ETS)
RORg(NR)
Egr2(Zf)
ETS1(ETS)
OCT:OCT(POU,Homeobox)
Etv2(ETS)
Rfx2(HTH)
Maz(Zf)
NFkB-p50,p52(RHD)
RFX(HTH)
ETV1(ETS)
OCT:OCT(POU,Homeobox)
0 10 20
–Log10(P-value)
Significantly enriched (P-value <0.05)
Not enriched
Target sequences with motif (%):
100%
10%
≤1%
B
Cluster 3 (6453 peaks)
BMLF1 signature
Promoter & Intragenic
(5054 peaks)
3461 genes
Top 5 peak-associated genes
LINC-PINT
FYN
PRKCH
GTDC1
CDC42SE2
–Log10(adjusted P-value)
0 2 4 6 8
T cell receptor signaling pathway
Cytokine signaling in immune system
Signaling by interleukins
Adaptive immune system
Programmed cell death
Signaling by Rho GTPases
Transcriptional regulation by RUNX3
Pos. reg. of cell cycle G1/S transition
Naïve
CM
EM
TEMRA
BMLF1
BRLF1
LMP2
(25)
(15)
(6)
IFNG
1 kb
C
Cluster 4 (7812 peaks)
LMP2 & CM signature
Promoter & Intragenic
(6328 peaks)
4339 genes
Top 5 peak-associated genes
RORA
BCL11B
TNFAIP8
NCK2
CDC14A
–Log10(adjusted P-value)
0 2 4 6
Innate immune system
Toll like receptor 3 cascade
MyD88 cascade initiated on PM
Th17 cell differentiation
Chemokine signaling pathway
Signaling by Rho GTPases
Toll like receptor 4 cascade
Signaling by interleukins
FoxO signaling pathway
T cell receptor signaling pathway
Naïve
CM
EM
TEMRA
BMLF1
BRLF1
LMP2
(44)
(17)
1 kb
IL7R
1 kb
D
Cluster 7 (5795 peaks)
BRLF1 signature
Promoter & Intragenic
(4508 peaks)
2849 genes
Top 5 peak-associated genes
IKZF2
HIVEP3
LOC105370401
RIN3
GAS7
–Log10(adjusted P-value)
0 2 4 6
Innate immune system
Chemokine signaling pathway
Natural killer cell mediated cytotoxicity
Fc receptor signaling pathway
RHO GTPase cycle
Signaling by NOTCH1
Signaling by VEGF
GH synthesis, secretion & action
IL2 family signaling
Positive reg. of cytokine production
Naïve
CM
EM
TEMRA
BMLF1
BRLF1
LMP2
(11)
(27)
1 kb
LYN
1 kb

**Supplementary Figure 4 (related to Figure 2): ATAC-seq chromatin accessibilities suggest different T cell traits of antigen-specific T cell populations. A**. HOMER transcription factor enrichments in K-means clusters from Fig. 2B. The 20 most significant TF enrichments are depicted, and the corresponding TF family are shown in parenthesis. **B-D**. Analyses of K-means cluster peaks showing the top 5 peak-associated genes (left) as defined by the largest number of promoter and intergenic peaks per gene, as well as the pathway and annotation enrichment results of promoter and intergenic peak-associated genes (middle). Selected ATAC-seq peak plots for each K-means cluster are shown (right).

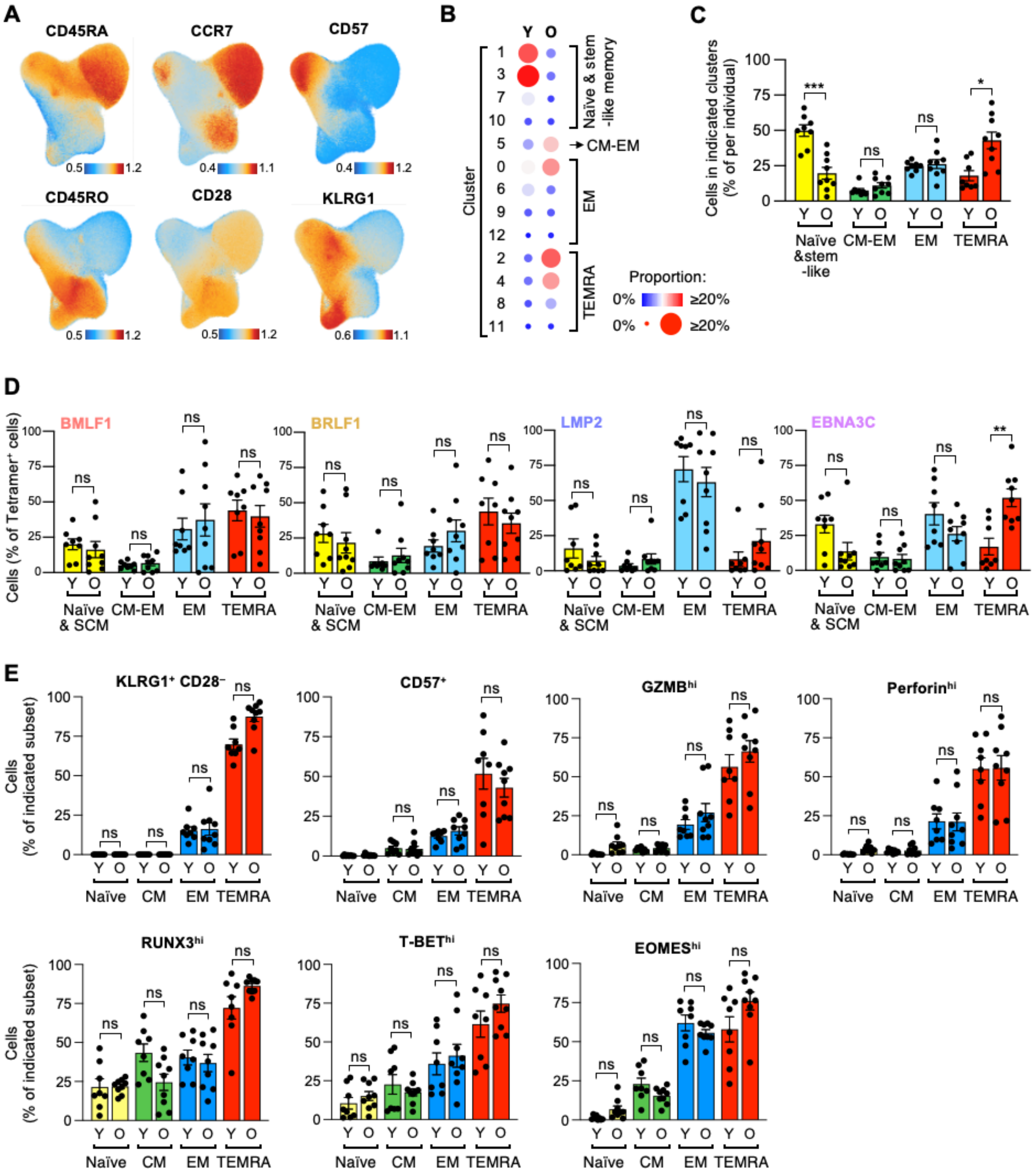
A
CD45RA
CCR7
CD57
CD45RO
CD28
KLRG1
B
Y O
Cluster
Naïve & stem -like memory
CM-EM
EM
TEMRA
Proportion:
0% ≥20%
C
Cells in indicated clusters (% of per individual)
Naïve &stem -like
CM-EM
EM
TEMRA
***
*
ns
D
Cells (% of Tetramer+ cells)
BMLF1
BRLF1
LMP2
EBNA3C
Naïve & SCM
CM-EM
EM
TEMRA
**
ns
E
Cells (% of indicated subset)
KLRG1+ CD28−
CD57+
GZMBhi
Perforinhi
RUNX3hi
T-BEThi
EOMEShi
Naïve
CM
EM
TEMRA
Y O
ns

**Supplementary Figure 5 (related to Figure 3): Aging-associated acquisition of T cell end-differentiation traits is dependent on antigen specificity. A**. Protein levels of selected subset markers in flow cytometry-based reference mapping. The corresponding reference map of bulk $CD8^+$ T cells is shown in Fig. 3A. **B-C**. Frequencies of bulk $CD8^+$ T cells of young (Y) and older (O) adults in reference map clusters shown in Fig. 3A are shown as bubble plots (B) or summarized as bar plots of classical T cell subsets (C). **D**. Distribution of classical T cell subset phenotypes of EBV-specific T cells based on reference mapping analyses in Fig. 3C. **E**. Flow cytometry quantification of indicated cell surface marker and intracellular proteins indicative of advanced differentiation in bulk $CD8^+$ T cell subsets. Data show the median (B) or mean ± SEM (C-E). All datapoints represent distinct biological replicates. Values O adults were compared to those of Y adults described in Fig. 1 and Supplementary Fig. 1. Data were compared by two-way ANOVA with Šídák's multiple comparisons test (C-E). *P<0.05, **P<0.01, ***P<0.001. ns, not significant.

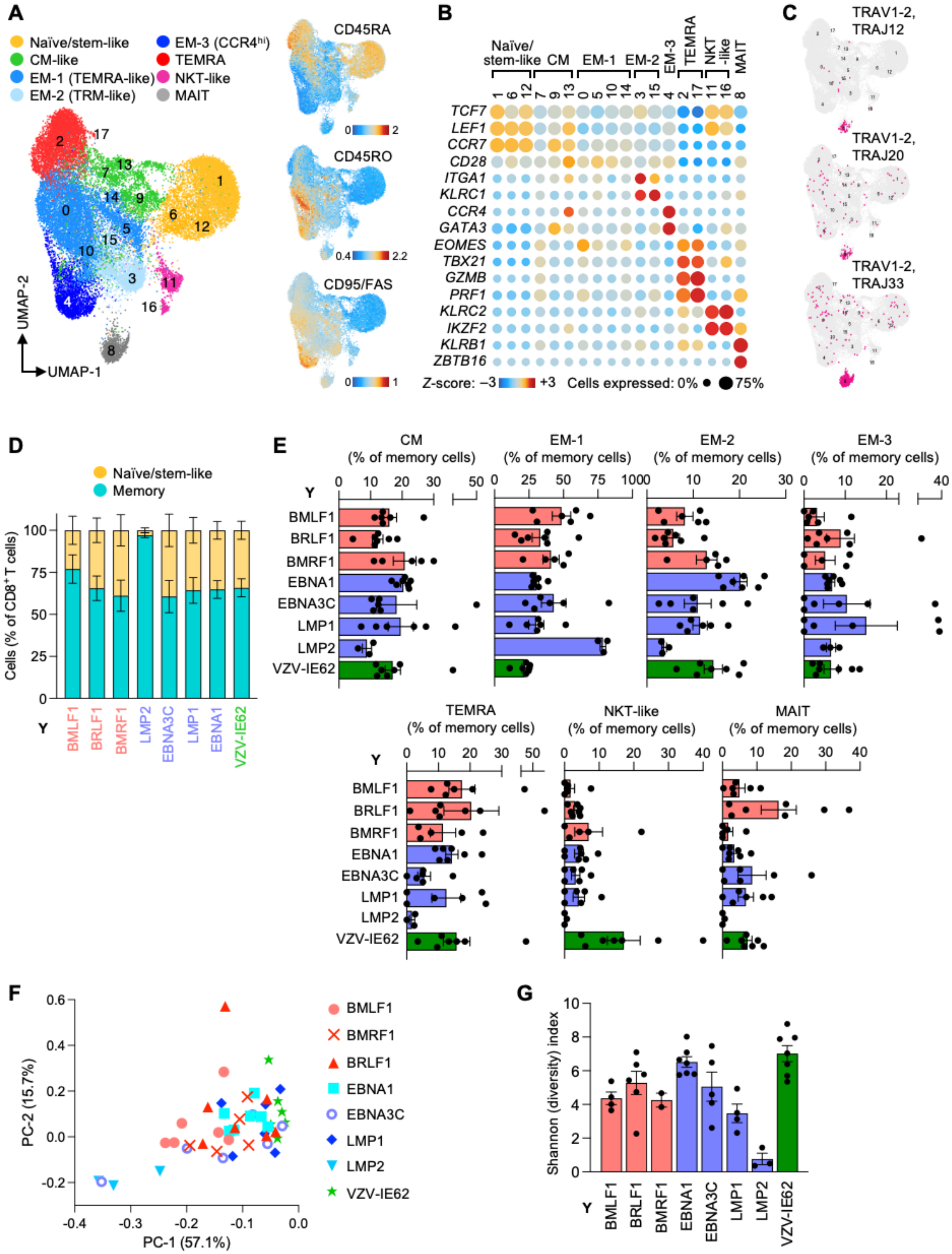

A
Naïve/stem-like
CM-like
EM-1 (TEMRA-like)
EM-2 (TRM-like)
EM-3 (CCR4hi)
TEMRA
NKT-like
MAIT
UMAP-2
UMAP-1
CD45RA
CD45RO
CD95/FAS
B
Naïve/ stem-like
CM
EM-1
EM-2
EM-3
TEMRA
NKT -like
MAIT
TCF7
LEF1
CCR7
CD28
ITGA1
KLRC1
CCR4
GATA3
EOMES
TBX21
GZMB
PRF1
KLRC2
IKZF2
KLRB1
ZBTB16
Z-score: −3 +3
Cells expressed: 0% 75%
C
TRAV1-2, TRAJ12
TRAV1-2, TRAJ20
TRAV1-2, TRAJ33
D
Naïve/stem-like
Memory
Cells (% of CD8+ T cells)
Y
BMLF1
BRLF1
BMRF1
LMP2
EBNA3C
LMP1
EBNA1
VZV-IE62
E
CM (% of memory cells)
EM-1 (% of memory cells)
EM-2 (% of memory cells)
EM-3 (% of memory cells)
TEMRA (% of memory cells)
NKT-like (% of memory cells)
MAIT (% of memory cells)
Y
BMLF1
BRLF1
BMRF1
EBNA1
EBNA3C
LMP1
LMP2
VZV-IE62
F
PC-2 (15.7%)
PC-1 (57.1%)
BMLF1
BMRF1
BRLF1
EBNA1
EBNA3C
LMP1
LMP2
VZV-IE62
G
Shannon (diversity) index
Y
BMLF1
BRLF1
BMRF1
EBNA1
EBNA3C
LMP1
LMP2
VZV-IE62

**Supplementary Figure 6 (related to Figure 4): Phenotypic diversity of antigen-specific CD8$^+$ T cells in single cell-sequencing analyses. A**. UMAP and KNN clustering of EBV-specific CD8$^+$ T cell sequencing data of young (Y) and older adults (left). Cell surface protein levels of 3 selected markers as assessed by CITE antibodies in single cell-sequencing data (right). **B**. Heatmap on RNA expression data across clusters defined in (A) depicting T cell subset and state markers. **C**. Expression of three T cell receptor (TCR) segments that are typical for MAIT cells. **D**. Proportion of naïve/stem-like cells and memory cells within antigen-specific T cells as phenotypically defined in (A-B). **E**. Quantitation of antigen-specific memory T cells across classical memory subsets as defined in (A-B). **F**. PCA plot based on the subset distribution frequencies of antigen-specific T cell samples in single cell sequencing clusters. **G**. T cell diversity as measured by the Shannon index. Only cells with TCR$\alpha$ and TCR$\beta$ sequences in single cell-sequencing data were used for the calculation. Data show the mean ± SEM (E, G). All datapoints represent distinct biological replicates.

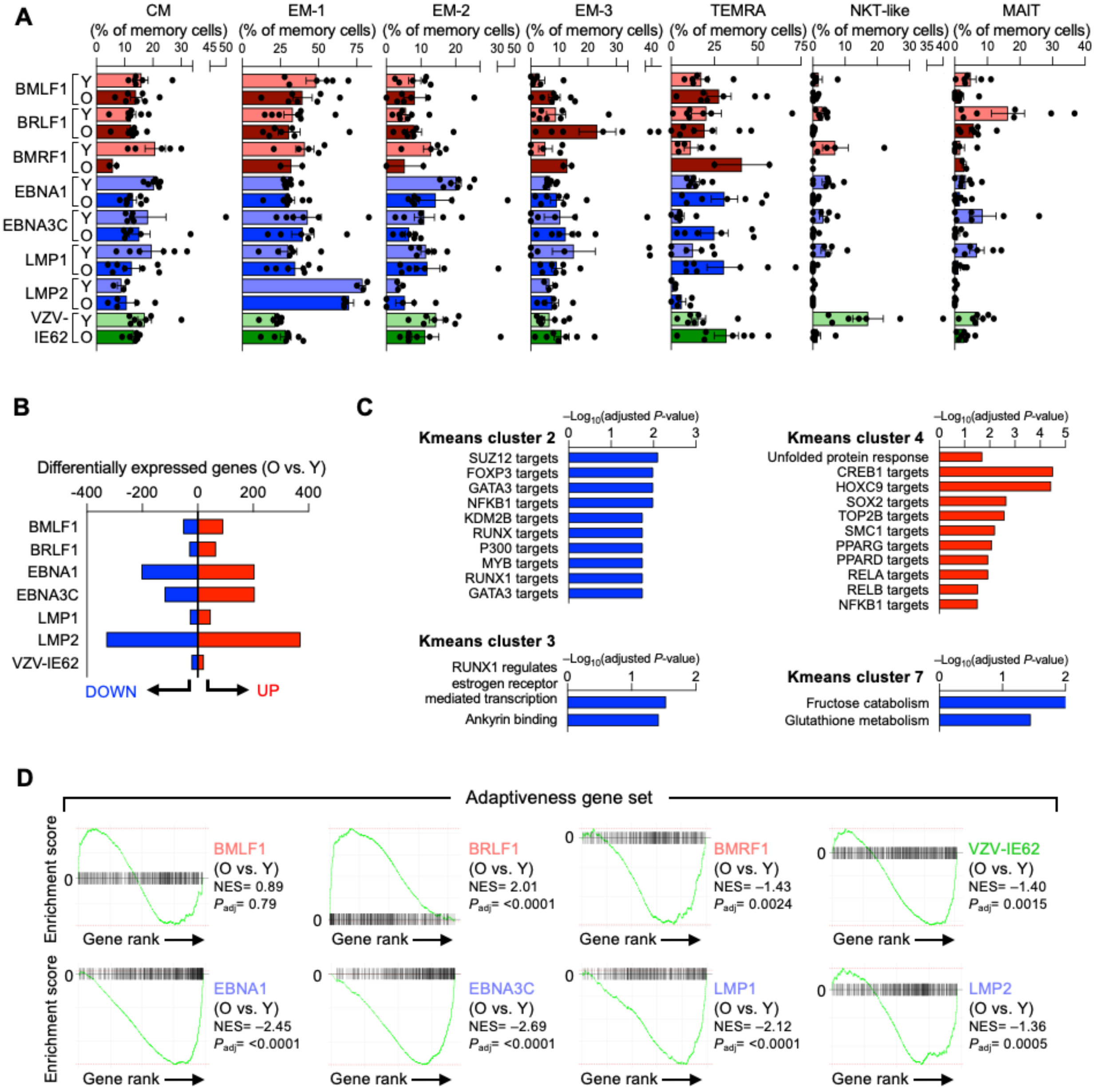


**E**

| | | BMLF1 | BRLF1 | BMRF1 | EBNA1 | EBNA3C | LMP1 | LMP2 | VZV-IE62 |
|---|---|---|---|---|---|---|---|---|---|
| Exhausted-vs-Effector (GSE9650) | NES | 1.35 | 1.28 | 0.83 | 1.33 | 1.37 | 1.77 | 1.4 | 1.61 |
| | *Padj* | 0.13 | 0.33 | 0.77 | 0.24 | 0.21 | 0.02 | 0.18 | 0.01 |
| Exhausted-vs-Memory (GSE41867) | NES | -0.83 | -0.76 | 0.98 | -0.86 | -0.75 | -0.79 | 0.74 | -0.81 |
| | *Padj* | 0.91 | 0.98 | 0.64 | 0.88 | 0.97 | 0.92 | 0.96 | 0.96 |
| Cellular senescence (GO-BP) | NES | 1.46 | 0.8 | -0.99 | 1.07 | 0.97 | 1.32 | -0.91 | 0.86 |
| | *Padj* | 0.12 | 0.98 | 0.64 | 0.66 | 0.73 | 0.32 | 0.96 | 0.96 |
| Cellular senescence (Reactome) | NES | -0.85 | 0.76 | -0.95 | 0.81 | 0.85 | 1.03 | -0.78 | -0.72 |
| | *Padj* | 0.91 | 0.98 | 0.64 | 0.88 | 0.91 | 0.48 | 0.96 | 0.96 |
| Cellular senescence (SenSig) | NES | 1.55 | 1.3 | 1.26 | 1.4 | 1.15 | 1.21 | 1.02 | 1.76 |
| | *Padj* | 0.12 | 0.33 | 0.64 | 0.24 | 0.56 | 0.42 | 0.9 | 0.01 |
| Senescence-associated secretory phenotype (SenMayo) | NES | 1.17 | 1.13 | 0.98 | -1 | -1.3 | 1.06 | 1.22 | 1.1 |
| | *Padj* | 0.22 | 0.34 | 0.64 | 0.7 | 0.06 | 0.48 | 0.18 | 0.32 |

**Supplementary Figure 7 (related to Figures 5 and 6): EBV-specific memory T cells do not acquire markers of exhaustion or cellular senescence with aging but lose adaptiveness signatures. A**. Quantitation of antigen-specific memory T cells across classical memory subsets as defined in Supplementary Fig. 6A comparing cells contrasting older (O) adults with cells of younger (Y) described in Supplementary Fig. 6A. **B**. Number of differentially expressed genes in each antigen-specific T cell population comparing Y and O samples. Differential expression analyses were based on pseudobulk comparison including only antigen-specific T cells having a memory phenotype. **C**. Pathway and annotation enrichment of genes in K-means clusters from Fig. 6C. **D**. Gene set enrichment analysis of DEGs in pseudobulk comparison for each antigen specificity. The "Adaptiveness" gene set was initially collated by Gutierrez-Arcelus *et al*.[36]. NES, normalized enrichment score. **E**. Gene set enrichment analysis of DEGs for T cell exhaustion-related gene sets or sets of genes expressed with cellular senescence. Data show the mean ± SEM in (A). All datapoints represent distinct biological replicates.
